# Simplifying principles underlying the highly complex peptide motif of the promiscuous chicken MHC class I molecule, BF2*21:01

**DOI:** 10.64898/2026.02.20.707067

**Authors:** Michael Harrison, Paul E. Chappell, Samer Halabi, Maria Danysz, Enock M. Mararo, Łukasz Magiera, Clemens Hermann, Michael J. Deery, Kathryn S. Lilley, Hans-Joachim Wallny, David W. Avila, William Mwangi, Venugopal Nair, Susan M. Lea, Nicola Ternette, Jim Kaufman

**Affiliations:** University of Cambridge, Department of Pathology, Tennis Court Road, Cambridge, CB2 1QP, U. K; University of Oxford, Sir William Dunn School of Pathology, South Parks Road, Oxford, OX1 3RE, U. K; University of Edinburgh, Institute for Immunology and Infection Research, Charlotte Auerbach Road, Edinburgh, EH9 3FL, U. K; University of Cambridge, Department of Biochemistry, Cambridge Centre for Proteomics, Tennis Court Road, Cambridge, CB2 1QR, U. K; Basel Institute for Immunology, Grenzacherstrasse 487, Basel, CH-4005, Switzerland; Institute for Animal Health, High Street, Compton, RG20 7NN, U. K; Pirbright Institute, Ash Road, Pirbright, GU24 0NF, U. K; University of Oxford, Department of Biology, South Parks Road, Oxford, OX1 3EL, U. K; St Jude Children’s Research Hospital, Department of Structural Biology, 262 Danny Thomas Place, Memphis TN 38105, U. S. A; University of Oxford, Centre for Immuno-Oncology, Old Road Campus, Headington, OX3 7DQ, U. K; University of Oxford, Jenner Institute, Old Road Campus, Headington, OX3 7FZ, U. K; University of Dundee, School of Life Sciences, Dundee, DD1 4HN, U. K; University of Cambridge, Department of Veterinary Medicine, Madingley Road, Cambridge, CB3 0ES, U. K; Yale University, Department of Immunobiology, 300 Cedar Street, New Haven CT 06520, U. S. A

## Abstract

In chickens and humans, classical class I molecules of the major histocompatibility complex (MHC) have a spectrum of correlated properties, including cell surface expression and peptide repertoire. Understanding the impact of these promiscuous generalists and fastidious specialists is of considerable interest. The promiscuous BF2 molecule from the chicken B21 haplotype, BF2*21:01, binds a wide range of peptides by remodelling the peptide-binding site, allowing co-variation of the anchor residues at peptide positions P_2_ and P_c-2_, and binding of an anchor residue at Pc. Using in vitro refolding assays with peptides and peptide libraries, determining thermostability and crystal structures, and analysing a chicken B21 cell line by immunopeptidomics, we found that BF2*21:01 will accommodate many possible combinations at P_2_ and P_c-2_, as well as several hydrophobic amino acids at P_c_. However, marked preferences for particular peptide lengths, particular amino acids at the three anchor residues, and particular combinations of amino acids at P_2_ and P_c-2_ as well as at P_3_ and P_c-3_ together lead to high frequencies of major peptides while still allowing the possibility of presenting a wide peptide repertoire. These simplifying principles may eventually allow predictions of pathogen peptides with stable binding for this iconic promiscuous class I molecule, as well as providing the data for more sophisticated peptide prediction methods.

## Introduction

The major histocompatibility complex (MHC) is a genetic region defined by the presence of a few highly polymorphic genes encoding classical class I and class II molecules, originally identified as transplantation antigens but now known to play crucial roles in the immune response to pathogens and tumours (Djaoud and Parham, 2020). The human MHC is an enormous genomic region with much recombination among hundreds of genes with a vast array of functions, with multigene families of classical class I and class II genes, strongly associated with autoimmunity and allergies but not so strongly with infectious pathogens (Trowsdale and Knight, 2013). In contrast, the chicken MHC is much smaller and simpler, with single dominantly-expressed class I and class II genes, and strong associations of MHC haplotypes with resistance and susceptibility to certain economically-important pathogens (Kaufman 2018; Miller et al 2016; Tregaskes and Kaufman, 2021).

In trying to understand the strong associations of the chicken MHC haplotypes with infectious disease, we discovered a hierarchy of alleles of the dominantly-expressed class I molecule, BF2. The original insight came from flow cytometry: erythrocytes from the B21 haplotype conferring resistance to the oncogenic herpesvirus that causes Marek’s disease have ten-fold less class I on the cell surface than from the susceptible B4, B15, B12 and B19 haplotypes (Kaufman et al., 1995). Since then, a suite of properties was found to correlate with this hierarchy, the most important being the peptide repertoire, defined as the range of different peptide sequences bound. An inverse correlation between the level of cell surface expression and the breadth of peptide binding was found, with less well-expressed alleles that have wide peptide repertoires correlating with resistance to Marek’s disease and other economically-important pathogens (Chappell et al., 2015; Kaufman, 2015; Kaufman, 2018; Tregaskes and Kaufman, 2021).

A similar inverse correlation between predicted peptide repertoire and cell surface expression was found for some human class I molecules, but a disease association that was the opposite: well-expressed “elite controller” alleles with narrow peptide repertoires correlated with slow progression from HIV infection to AIDS (Chappell et al., 2015). This finding led to the hypothesis of generalist and specialist class I alleles, with low-expressing promiscuous alleles generally protecting from many pathogens while high-expressing fastidious alleles bound special peptides to protect against particular pathogens (Chappell et al., 2015; Kaufman, 2018). A hierarchy of tapasin-dependence and ease of in vitro refolding for certain human class I alleles (Rizvi et al., 2014) was found to correlate with cell surface expression and predicted peptide repertoire (Kaufman, 2015; Kaufman, 2018; Tregaskes and Kaufman, 2021) and eventually shown to associate with slow progression once elite controllers were removed from analysis (Bashirova et al., 2020), thus supporting the generalist-specialist hypothesis. In addition, a hierarchy of human class II alleles based on predicted peptide repertoire was reported (Manczinger et al., 2019).

There are presumably various biochemical mechanisms leading to this hierarchy of class I alleles, including specificity of peptide transport by TAPs in chickens (Tregaskes et al., 2016; Walker et al., 2011), dependence of class I alleles on tapasin (and perhaps TAPBPR) in humans (Bashirova et al., 2020; Kaufman, 2015; Kaufman, 2018; Rizvi et al., 2014), and specificity of peptide binding by the class I alleles themselves (Chappell et al., 2015; Han et al., 2023; Koch et al., 2008; Paul et al., 2013; Shaw et al., 2007; Wallny et al., 2006; Xiao et al., 2018; Zhang et al., 2012). Several modes of promiscuous binding have been reported for dominantly-expressed chicken class I molecules (Halabi and Kaufman, 2022).

Despite requiring three anchor residues, BF2*21:01 achieves a wide peptide repertoire by remodelling the peptide-binding site in a previously unprecedented manner that involves co-variation between the amino acids at positions P_2_ and P_c-2_, as shown by structures bearing six peptides with very divergent sequences (Chappell et al., 2015; Koch et al., 2008). Since this iconic chicken class I molecule is so frequent across the world and so important for resistance to economically-important viral diseases (Kaufman, 2018), we wished to better understand the rules by which peptides bind to BF2*21:01, starting with refolding the class I molecule with different peptides and peptide libraries, determining thermostabilities, determining structures and creating models to illustrate some of the results, and finishing with immunopeptidomics of class I molecules from a B21 cell line.

## Results

### The class I molecule(s) from the B21 haplotype bind a wide variety of peptides

Gas phase sequencing of peptides eluted from B21 class I molecules and separated by HPLC gave a wide range of sizes and sequences with no obvious anchor residues (Fig. 1), in stark contrast to the class I molecules from B4, B12, B15 and B19 which gave mostly octamers with clear peptide motifs (Kaufman et al., 1995; Wallny et al., 2006). The heterogeneity in length of peptides from blood and spleen cells is likely to have been due to proteolysis, since the single preparation from a B21 chicken cell line gave only 10mers and 11mers, including TNPESKVFYL from which the octamer PESKVFYL from blood samples (that failed to refold in vitro) was apparently derived. The wide range of sequences were eventually understood to mean that BF2*21:01 molecules would accommodate a variety of anchor residues at three positions, with co-variation between the anchor residues at P_2_ and P_c-2_ (position 2 and the position two before the C-terminus of the peptide), as found by refolding and crystallography (Chappell et al., 2015; Koch et al., 2008).

**Figure 1.**
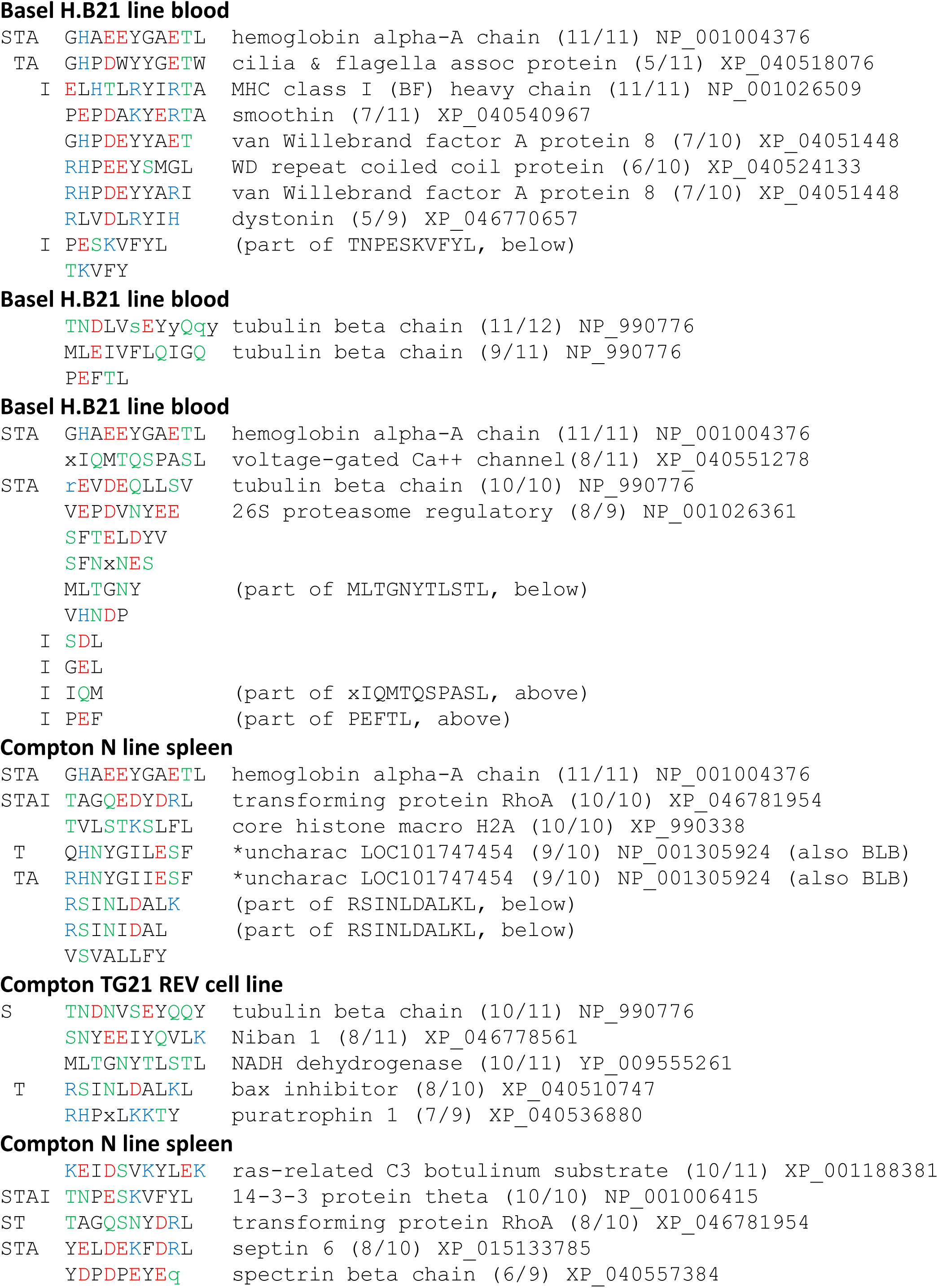
Peptides isolated from class I molecules of chicken blood, spleen and a B21 cell line separated by HPLC with single peaks subjected to gas phase sequencing show a range of lengths and sequences. Methods are detailed in Chappell et al., 2015; Koch et al., 2008; Wallny et al., 2006. The peptides are grouped by the six experiments with locations and cell sources indicated. Sequences are in single letter amino acid code, small letter indicates dominant amino acid in a position, and x indicates ambiguous call. The amino acids are coloured throughout this paper according to their primary characteristic (with the understanding that some amino acids could be characterised in more than one way, eg. Y both hydrophobic and polar, K basic and hydrophobic, etc): red, acidic, D and E; blue, basic, H, K and R; green, polar, N, Q, S and T; black, hydrophobic, A, C, F, G, I, L, M, P, V, W and Y). Designations on the left indicate synthetic peptides used for structures (S), thermostability assays (T) or assembly assays (A), or found as natural peptide by immunopeptidomics (I). Chicken genes from which a (partial) peptide sequence could be identified are presented, with the number of amino acids positions matching compared to total length in brackets.

To better understand the requirements of peptide binding to BF2*21:01, refolding in vitro followed by size exclusion chromatography (hereafter called “assembly”) was carried out with individual peptides. Four outcomes were routinely found: no monomer peak reflecting no assembly, a relatively sharp monomer peak reflecting a stable binding, a broader monomer peak that reflected monomers falling apart during chromatography and a sharp peak at the position of BF2*21:01 heavy chain alone that presumably reflected monomers that fell apart before or during chromatography (Fig. 2).

**Figure 2.**
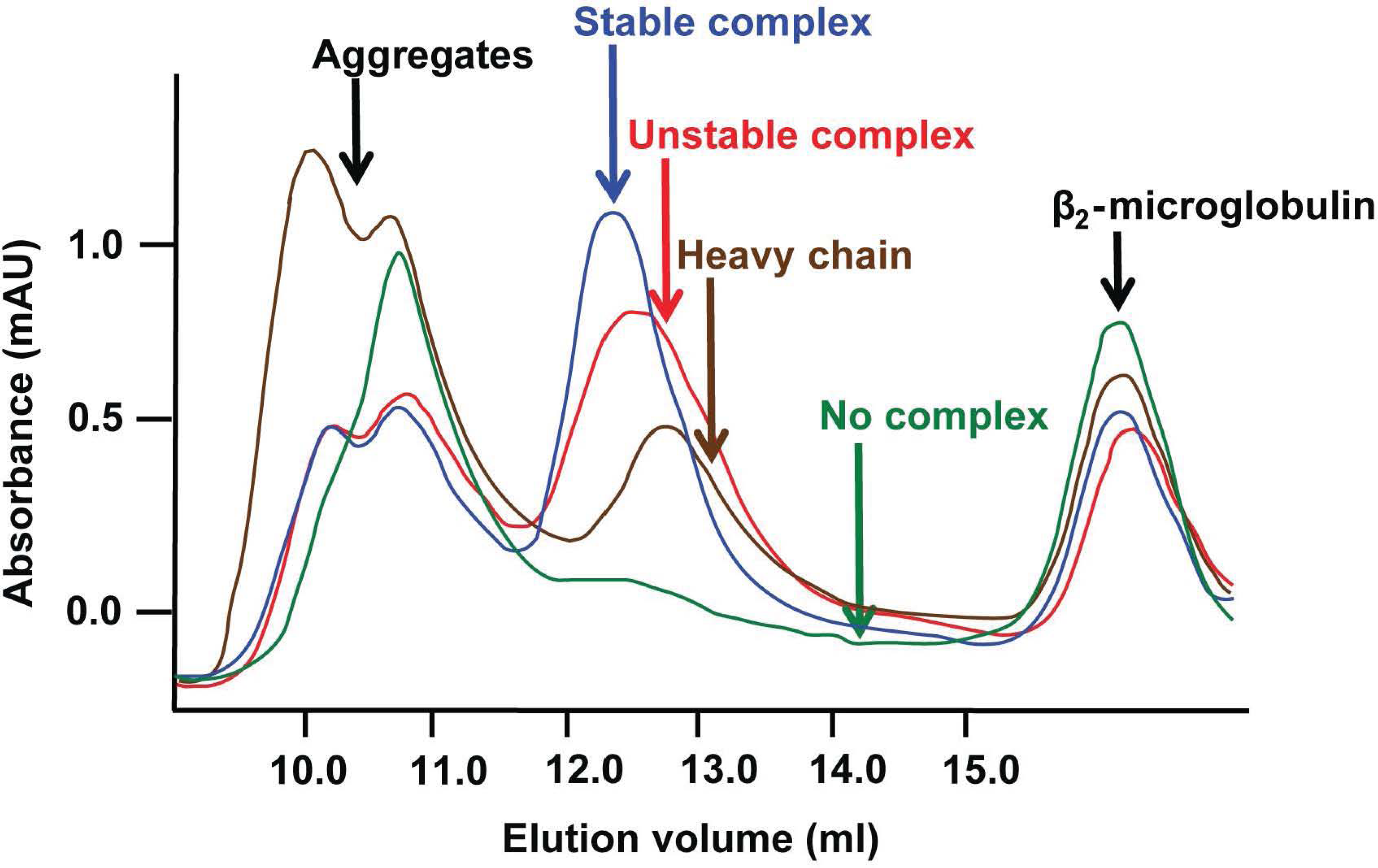
Representative SEC trace for assembly assays show four typical outcomes for BF2*21:01. After refolding of bacterially-expressed heavy chain and β_2_-microglobulin with synthetic peptides according to published methods (Chappell et al., 2015; Koch et al., 2008), samples were spun in a cooled microfuge at full speed for 30 minutes, the supernatant was put through a 0.2 microfilter and then the flow-through was loaded on a SEC column (in this case, a Superdex 200 10/300 GL column) attached to an AKTA FPLC instrument. In every run, peaks at and slightly after the exclusion volume represent aggregated material, and a peak before the inclusion body represented refolded β_2_-microglobulin. In between, a sharp peak represented stably refolded monomers of heavy chain, β_2_-microglobulin and peptide (peptide with blue trace), a broader peak beginning at the same place as the monomer peak presumably represented refolded monomers that were dissociating during chromatography (red), a later broad was at the same position as heavy chain refolded alone (brown) and for some peptides no peak in between the aggregate peaks and the β_2_-microglobulin peak, representing no successful refolding (green). The x-axis represents elution volume in ml, while the y-axis represents OD_215_ in mAU.

Starting with two peptides for which the first crystal structures for BF2*21:01 were determined (Koch et al., 2008), assembly assays with single peptides substituted at positions P_2_, P_c-2_ and P_c_ showed that some (combinations of) amino acids were tolerated while others were not (Fig. 3). For the 11mer GHAEEYGAETL, the basic amino acid His at P_2_ allowed both acidic Asp and Glu at P_c-2_ but the hydrophobic Leu at P_2_ gave unstable binding. However, substituting basic Lys and Arg at P_2_ only allowed Asp at P_c-2_. The 10mer REVDEQLLSV allowed many hydrophobic amino acids (except Ala) at P_c-2_, but substituting acidic Asp at P_2_ allowed no stable peptide binding with the same amino acids at P_c-2_. Indeed, both the 11mer and the 10mer peptides with substitutions of hydrophobic amino acids at P_2_ failed to bind in a stable way. The 11mer substituted with a Glu at P_2_ alone or with Leu at P_c-2_ (as in the 10mer) only bound well when P_3_ was changed to Val (as in the 10mer). In contrast, the 10mer tolerated His at P_2_ and Glu at P_c-2_ (as in the 11mer), except with Ala at both P_3_ and P_c-3_ (as in the 10mer). Substitution at P_c_ in both the 10mer and 11mer showed that Leu, Phe and Val but not Ala or Trp were tolerated. Assembly assays with derivatives of the 11mer show that the 10mer and 11mer bound well, but 9mers and below were not as stable (Fig. 4). Overall, the rules for binding seemed complex.

**Figure 3.**
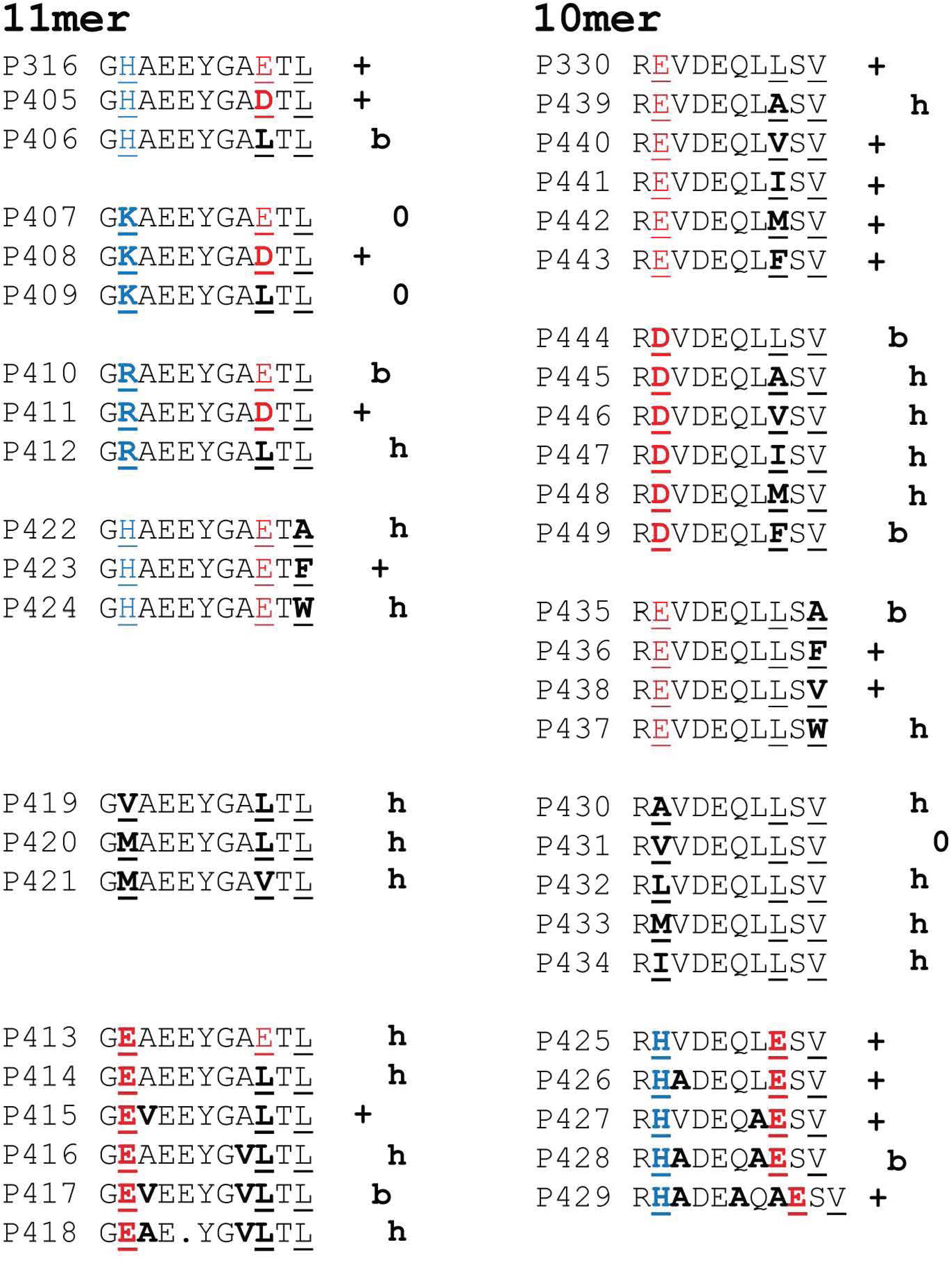
Assembly of BF2*21:01 molecules by refolding in vitro is markedly affected by single amino acid differences between peptides. Bacterial-expressed heavy chain and β_2_-microglobulin were refolded in vitro with single synthetic peptides (in-house identification of peptide batch by P and number), shown as single letter code with anchor residues underlined and coloured (red, acidic; blue, basic; black hydrophobic), changes from original peptide bolded, and a dot to indicate a residue skipped. Result after analysis by SEC given as +, sharp peak at position of monomer; b, broad peak starting at position of monomer; h, sharp peak at position of heavy chain alone; 0, no peak in between aggregated material at the exclusion volume of the column and the peak representing refolded β_2_-microglobulin. Methods as in Fig. 2.

**Figure 4.**
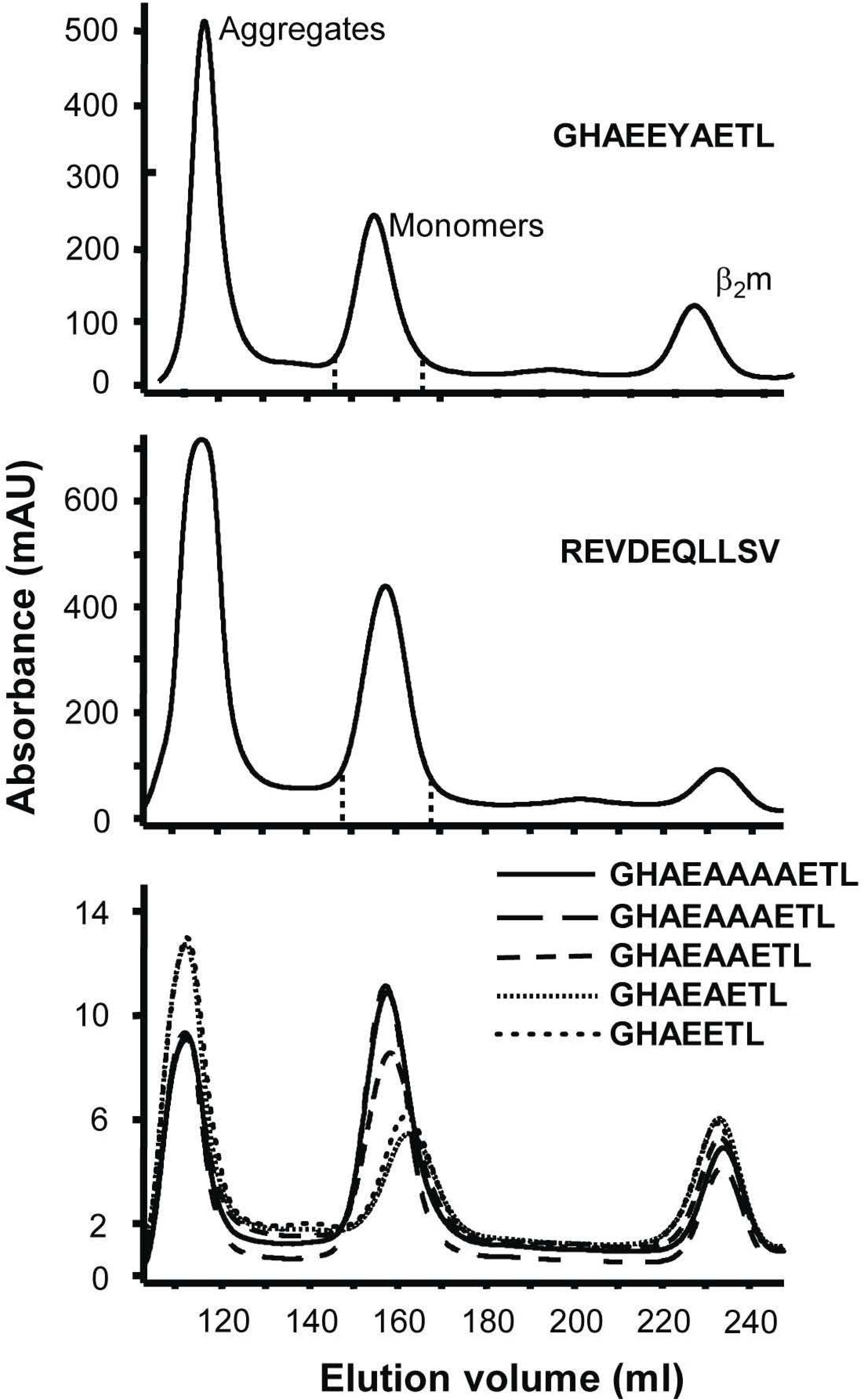
The original 10mer (REVDEQLLSV) and 11mer (GHAEEYGAETL) peptides refold with BF2*21:01 to give stable monomers as do 11mer and 10mer derivative peptides, but 9mer, 8mer or 7mer peptides give heavy chain only peaks. Assembly assays for representative 10mer and 11mer peptides give peaks for stable monomers of BF2*21:01, but shorter peptides based on the 11mer do not. Single synthetic peptides were refolded with bacterial-expressed heavy chain and β_2_-microglobulin with synthetic peptides, prepared for SEC as described in the legend for Fig. 2, and analysed in this case with a HiLoad 26/60 Superdex 200 column attached to an AKTA FPLC instrument. The peptides 11mer GHAEEYAETL (top panel), 10mer REVDEQLLSV middle panel), 11mer GHAEAAAAETL and 10mer GHAEAAAETL (bottom panel) gave sharp monomer peaks, while the 9mer GHAEAAETL, 8mer GHAEAETL and 7mer GHAEETL gave a delayed broad peak indicative of unstable binding or heavy chain.

Some of these diverse results could be understood by structures and models of the peptides bound to BF2*21:01. Structures with amino acid substitutions in the 11mer peptide GHAEEYGAETL bind with Asp at P_c-2_ and either His or Arg at P_2_ (5AD0 and 5ADZ), comparable to the original structure (3BEV) (Fig. 5A). Modelling the substituted peptide with Arg at P_2_ and Glu at P_c-2_ shows a steric clash that can explain why this peptide did not refold with BF2*21:01 (Fig. 5A). At P_c_, substitution of a Trp instead of the original Leu in this 11mer peptide leads to steric clashes with pocket F of BF2*21:01 (Fig. 5B).

**Figure 5.**
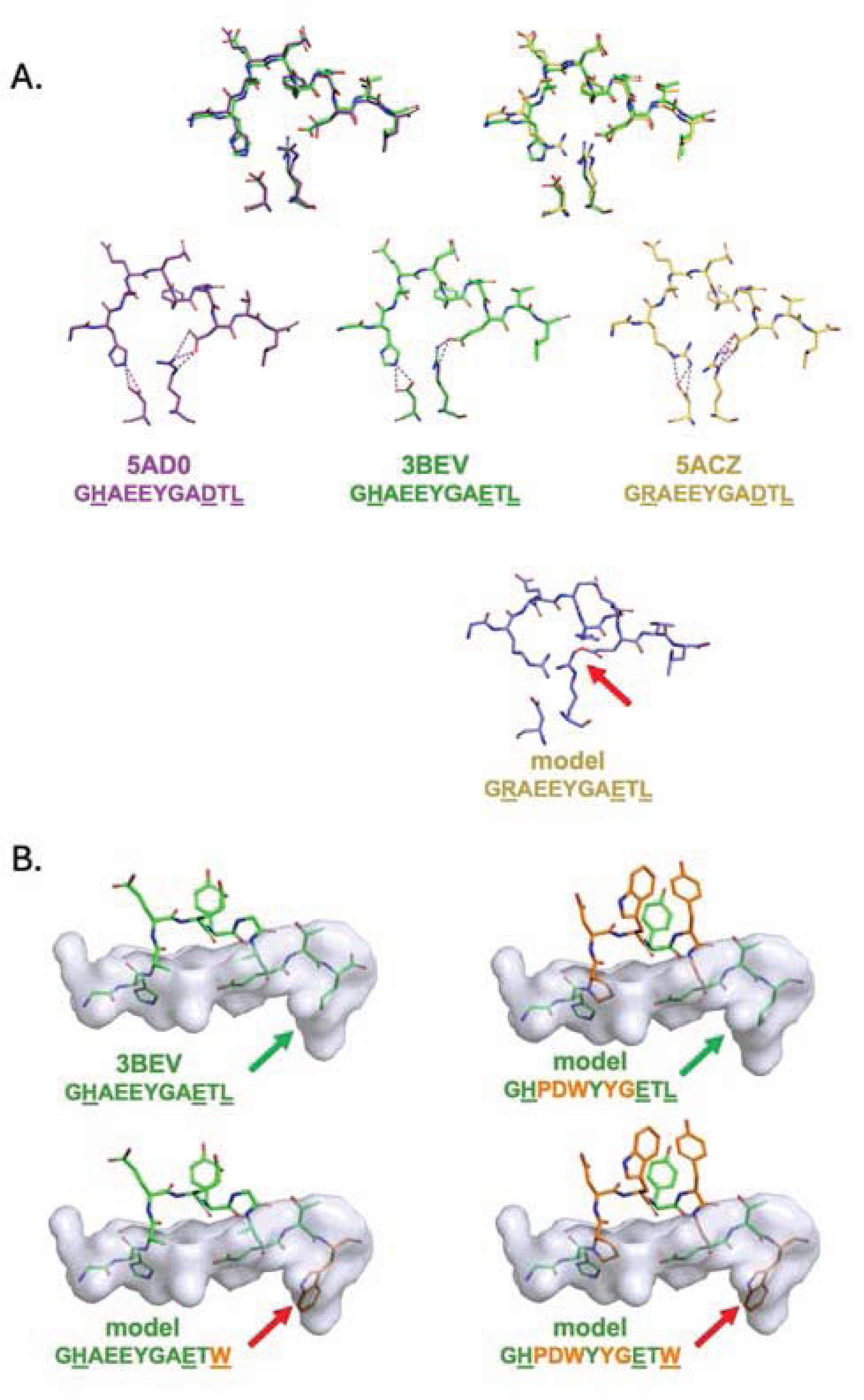
Crystal structures of BF2*21:01 with peptides that refold to give stable monomers and models of structures with peptides that either did not refold to give stable monomers or refolded to give monomers with low thermal stability. A. Structures were determined for G<u>H</u>AEEYGA<u>E</u>T<u>L</u> (3BEV), G<u>H</u>AEEYGA<u>D</u>T<u>L</u> (5AD0) and G<u>R</u>AEEYGA<u>D</u>T<u>L</u> (5ACZ), which all refolded successful to give stable monomers, while G<u>R</u>AEEYGA<u>E</u>T<u>L</u> did not (see Fig. 3), all models of which showed steric clashes (one depicted, red arrow). B. Two 11mers refolded to give monomers, which were much more stable for G<u>H</u>AEEYGA<u>E</u>T<u>L</u> than G<u>H</u>PDWYYG<u>E</u>T<u>W</u> (see Fig. 6). Models of G<u>H</u>AEEYGA<u>E</u>T<u>W</u> and G<u>H</u>PDWYYG<u>E</u>T<u>W</u> show steric clashes (red arrows), but the crystal structure with G<u>H</u>AEEYGA<u>E</u>T<u>L</u> (3BEV) and a model of G<u>H</u>PDWYYG<u>E</u>T<u>L</u> do not (green arrows). Underlined positions indicate anchor residues; bold letters indicate amino acid substitutions, modelled positions in orange. Methods for refolding and analysis as in Fig. 2, modelling done as detailed in Materials and Methods, structure from Koch et al., 2008.

The thermostability of BF2*21:01 molecules refolded with various peptides confirms and extends these findings (Fig. 6). From the peptides eluted from B21 cells, one 11mer and eight 10mers (of which the structures have been determined for five, Chappell et al., 2015; Koch et al., 2008) were tested. The 10mers REVDEQLLSV and TNPESKVFYL were the most stable and the 10mer RSINGLDALKL and the 11mer GHPDWYYGETW were the least stable. The 11mer GHAEEYGADTL (a derivative of the original GHAEEYGAETL) was of middling stability, but substitution of P_2_ with Arg or Lys reduced the stability to the same as the 11mer GHPDWYYGETW. Similarly, the high stability of the 10mer REVDEQLLSV was reduced by substitution of P_c-2_ to Val, Ile or Met, or by the combination of His at P_2_ and Glu at P_c-2_ (as in the 11mer GHAEEYGADTL). The substitution of Phe at P_c_ lowered the stability of REVDEQLLSV only slightly, but the substitution of Leu at P_c_ in GHPDWYYGETW raised the stability significantly, consonant with models of the two peptides bound to BF2*21:01 (Fig. 5B). Derivatives of the 11mer, GHAEAA(A,AA)ETL, and of the 10mer, RHADEAQAESV, showed that 10mers were more stable than either 9mers or 11mers (Fig. 6).

**Figure 6.**
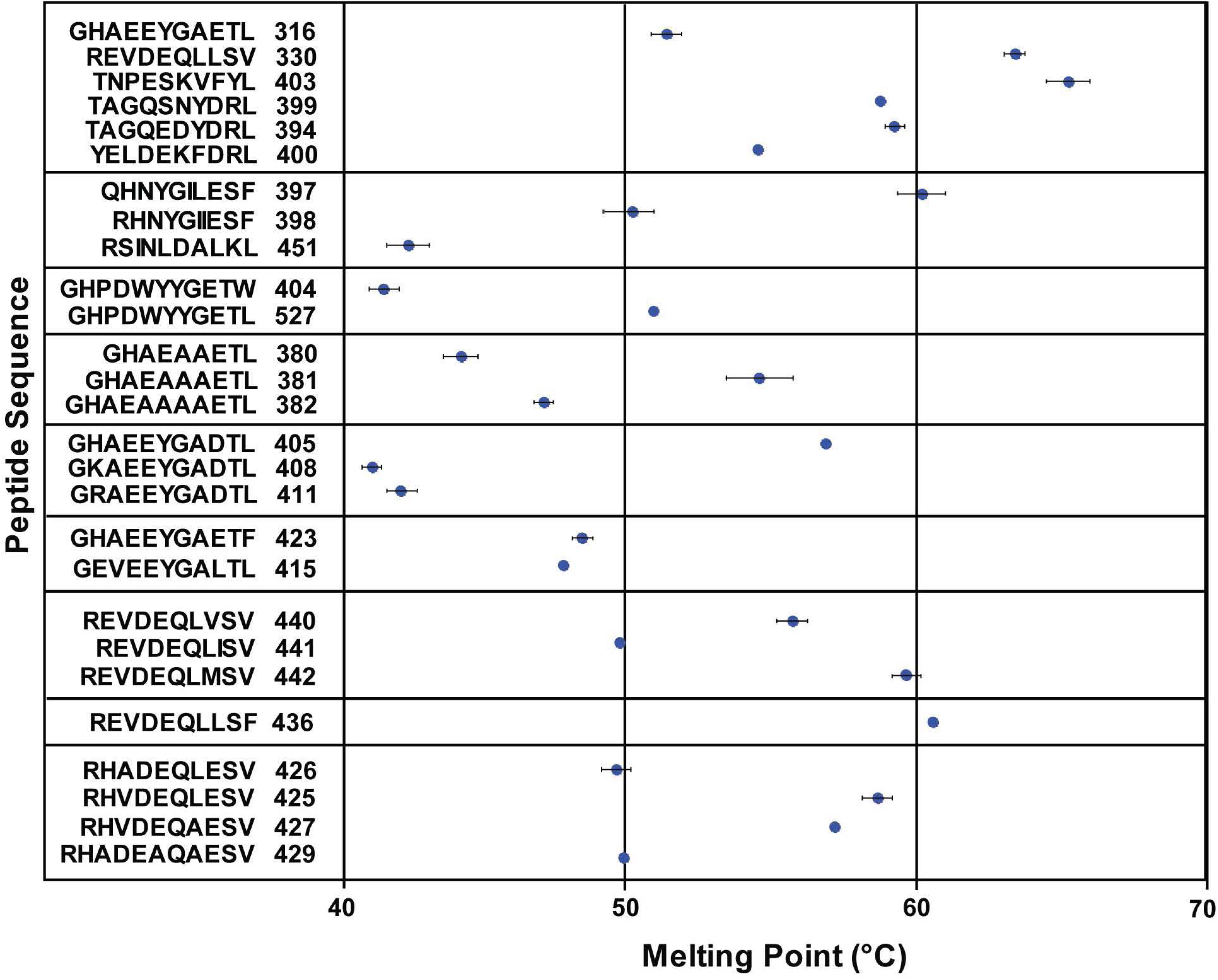
Peptide length and amino acids at peptide positions P_2_, P_3_, P_c-3_, P_c-2_ and P_c_ affect stability of BF2*21:01 molecules. Bacterial-expressed heavy chain and β_2_-microglobulin were refolded in vitro with various synthetic peptides and melting point measured as 50% uptake of a fluorescent dye that binds to hydrophobic regions, as described in Materials and Methods. Assays were performed in triplicate with error bars indicating standard errors. Peptide sequences are in single letter code with in-house identification of peptide batch by P and number.

### Assembly assays with peptide libraries show great variation in anchor and peptide backbone residues

To expand the range of amino acids examined, assembly assays followed by mass spectrometry were conducted with peptide libraries based on the 11mer and 10mer, starting with single-substitution libraries assessed by MALDI-TOF (Fig. 6). The libraries contained roughly equimolar amounts of all amino acids except Cys; given the molecular mass, Ile and Leu could not be distinguished, nor could Gln and Lys. Two concentrations of peptide were used, one at the usual 10-fold molarity over heavy chain and one at only 2-fold, with the idea that at 10-fold (but not 2-fold) a particularly high affinity peptide might be in sufficient concentration to compete out other binding peptides. In fact, there were few cases where the two concentrations gave much difference.

Using these single-substitution libraries based on the 11mer GHAEEYGAETL and the 10mer REVDEQLLSV (Fig. 7), substitution at P_2_ for the 11mer showed predominantly His with much lesser amounts of Arg, Ser, Gln/Lys and Asn, while for the 10mer both His and Glu were predominant with much lesser amounts of Asn, Asp, Ala and Ser. Substitution at P_c-2_ for the 10mers REVDEQLLSV and RDVDEQLLSV showed Phe followed by Ile/Leu, Met, Gln/Lys and Val. For both the 11mer and 10mer, substitutions at Pc showed Ile/Leu and Phe followed by Met and Val. Substitutions at P_3_ and P_c-3_ showed a variety of patterns.

**Figure 7.**
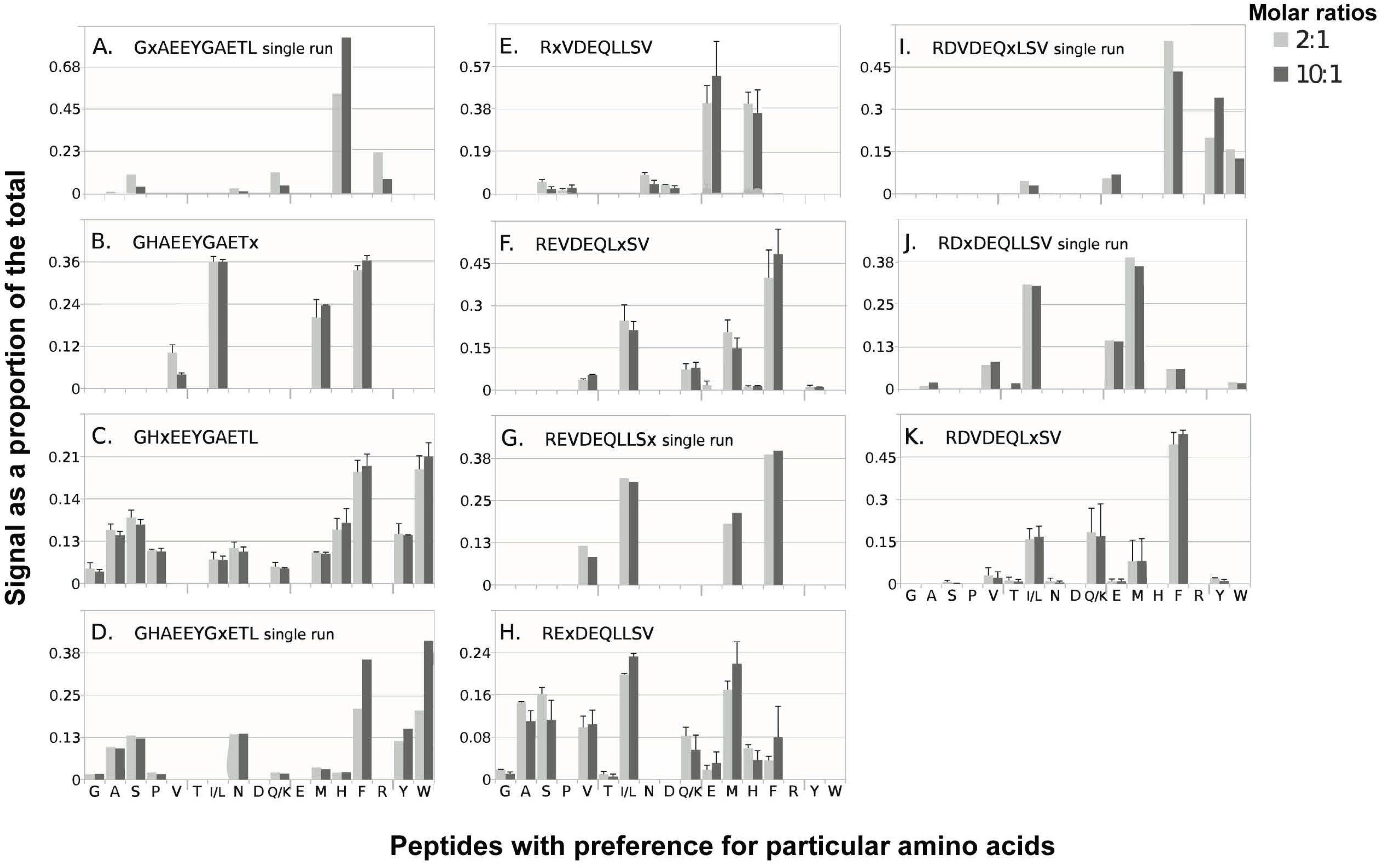
Assembly assays with single-substitution peptide libraries based on the 11mer GHAEEYAETL and the 10mer REVDEQLLSV show marked preferences for particular amino acids at P_2_, P_3_, P_c-3_, P_c-2_ and P_c_ positions binding to BF2*21:01. Bacterial-expressed heavy chain and β_2_-microglobulin were mixed with peptide libraries (19 amino acids, all except Cys, at one position) at two molar ratios (1:2:10 as usual, and 1:2:2 to control for strong binding peptides) and separated by SEC as described in the legend to Fig. 2; monomer peaks were collected, concentrated, treated with acid, lyophilised and analysed by MALDI-TOF. Bar charts show proportion of total signal (with error bars when present showing standard error around the mean for separate refolding experiments) on y-axis for mass/charge (m/z) positions corresponding to peptides with particular amino acids on the x-axis (single letter code; Ile and Leu were indistinguishable, as were Gln and Lys).

To begin to understand the co-variation between P_2_ and P_c-2_, double-substitution libraries were created for the 11mer GHAEEYGAETL and the 10mer REVDEQLLSV with the assembly assays analysed by MALDI-TOF (Fig. 8). This approach has serious limitations, not being able to distinguish between combinations that gave the same peaks (such as Arg-Ser and Asp-Lys) and or which position the amino acids were in (Asp at P_2_ and Ser at P_c-2_, or vice versa). A further issue was the temperature stability of the complexes, so after refolding at 4°C but before chromatography at room temperature, aliquots were incubated for 18 hours at 4°C (as for the refolding) and at 42°C (roughly the body temperature of a chicken), compared to the control with no incubation. The proportions of most combinations did not change significantly between the control and 42°C incubation, but some combinations increased in proportion (four out of 15 combinations for the 11mer, and four out of 37 combinations for the 10mer), suggesting that such combinations are more stable or have slower dissociation times than those combinations that reduced in proportion. Perhaps the most important finding was that far fewer combinations of P_2_ and P_c-2_ gave rise to stable peptide binding for the 11mer than for the 10mer, suggesting that this 11mer binds less stably and thus tolerates fewer changes than the 10mer.

**Figure 8.**
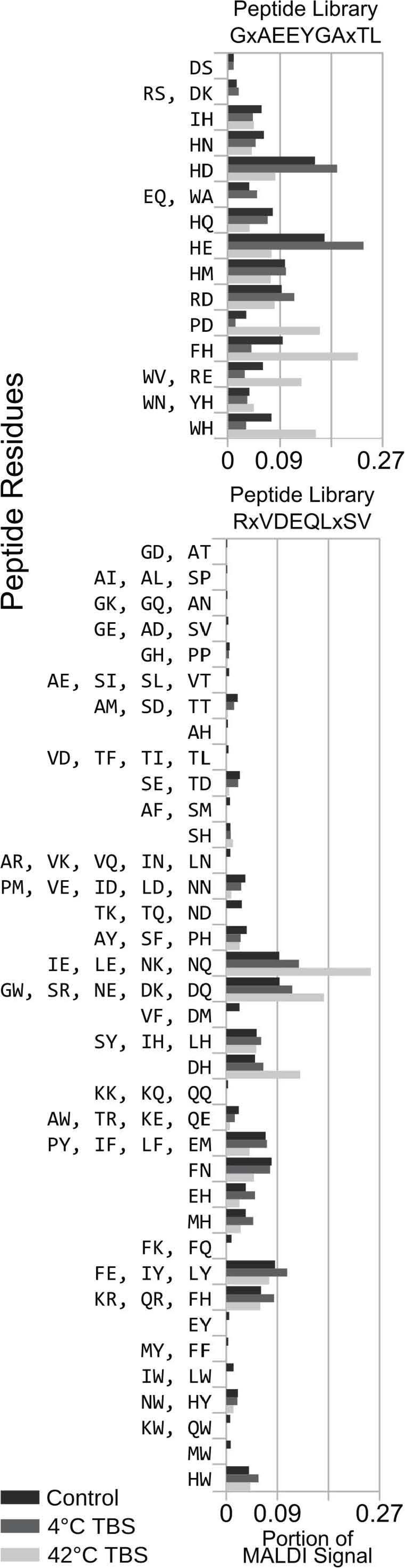
Assembly assays with double-substitution peptide libraries based on the 11mer GHAEEYAETL (top panel) and the 10mer REVDEQLLSV (bottom panel) show marked preferences for particular amino acids combinations at P_2_ and P_c-2_ positions binding to BF2*21:01, with more combinations found for the 10mer. Bacterial-expressed heavy chain and β_2_-microglobulin were mixed with peptide libraries (19 amino acids, all except Cys, at two positions indicated by x), separated immediately by SEC as described in the legend to Fig. 2, or incubated at 4°C or 42°C for 18 hours before SEC; monomer peaks were collected, concentrated, treated with acid, lyophilised and analysed by MALDI-TOF. Bar charts show proportion of total signal (with error bars when present showing standard error around the mean for separate refolding experiments) on x-axis for mass/charge (m/z) positions corresponding to peptides with particular amino acid combinations on the y-axis (single letter code; Ile and Leu, Gln and Lys, and amino acids present in P_2_ versus P_c-2_ positions were indistinguishable).

The examination of double-substitution libraries analysed by MALDI-TOF was expanded to other backbones (that is, sequences in which two positions were randomised): five other 11mer peptides (GHPDWYYGETW and four derivatives of the 11mer GHAEEYGAETL) and five 10mers (all from the original peptides found by gas phase sequencing, including the original REVDEQLLSV). For those libraries substituted at P_2_ and P_c-2_, four 11mers gave between 16 and 24 combinations, while the six 10mers gave between 32 and 55 combinations (Figs. 9, 10), so again the 10mers seemed to tolerate more combinations. Two other 11mer libraries examined substitutions at P_3_ and P_c-3_, which allowed many more combinations (Fig. 9).

**Figure 9.**
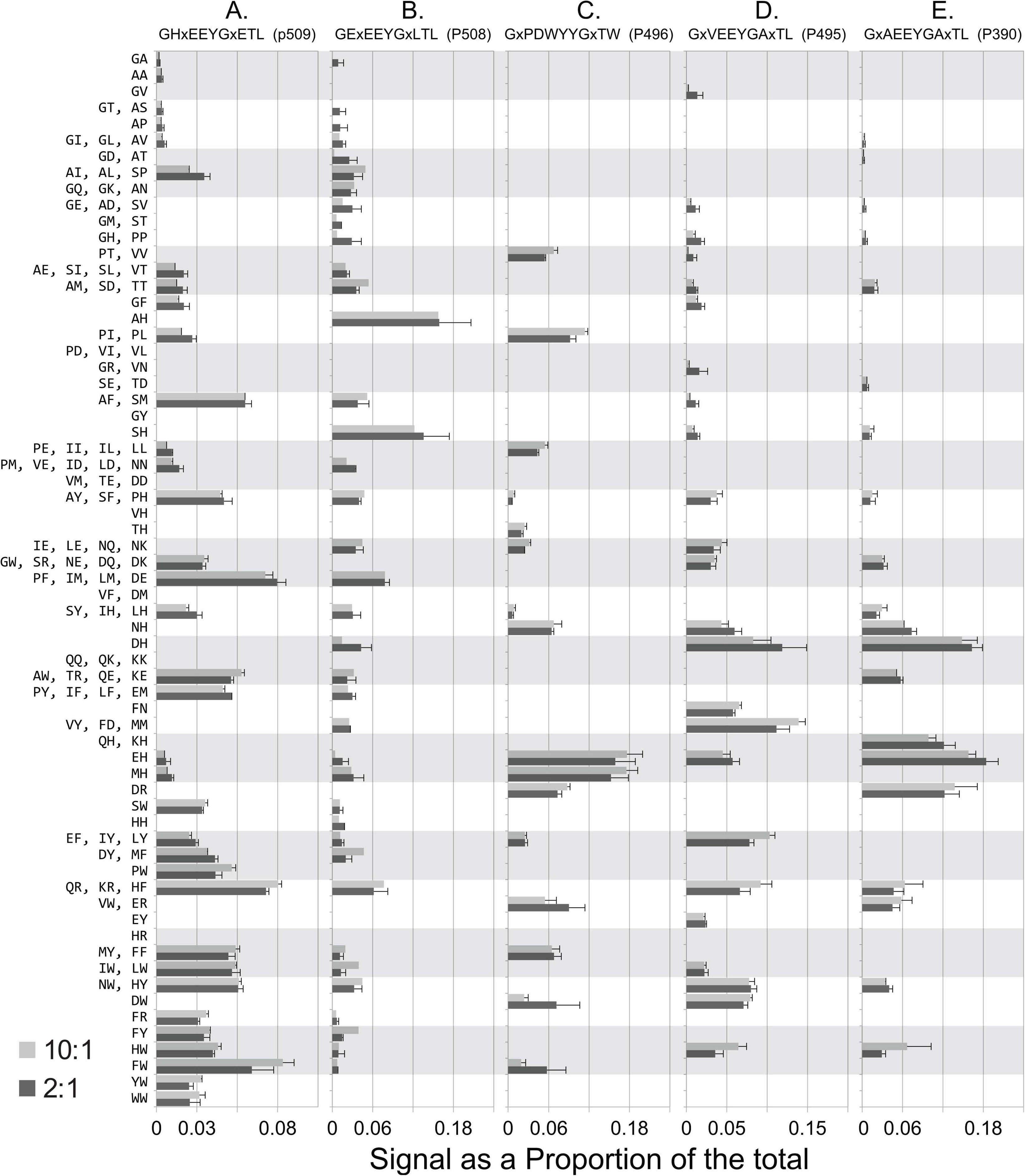
Panels A-E. Assembly assays with double-substitution peptide libraries based on five 11mer peptides (GHPDWYYGETW and four derivatives of the 11mer GHAEEYGAETL) and five 10mers (all from the original peptides found by gas phase sequencing, including the original REVDEQLLSV) show different preferences for particular amino acid combinations at P_2_, P_3_, P_c-3_ and P_c-2_ positions binding to BF2*21:01, with more combinations found for 10mer peptides. Bacterial-expressed heavy chain and β_2_-microglobulin were mixed with peptide libraries (19 amino acids, all except Cys, at one position, indicated by x in the peptide sequence) at two molar ratios (1:2:10 as usual, and 1:2:2 to control for strong binding peptides) and separated by SEC as described in the legend to Fig. 2 and separated immediately by SEC; monomer peaks were collected, concentrated, treated with acid, lyophilised and analysed by MALDI-TOF. Bar charts show proportion of total signal (with error bars when present showing standard error around the mean for separate refolding experiments) on x-axis for mass/charge (m/z) positions corresponding to peptides with particular amino acid combinations on the y-axis (single letter code; Ile and Leu, Gln and Lys, and amino acids present in P_2_ versus P_c-2_ positions were indistinguishable). Panels: A. GHxEEYGxETL, B. GExEEYGxLTL, C. GxPDWYYGxTW, D. GxVEEYGAxTL, E. GxAEEYGAxTL.

**Figure 10.**
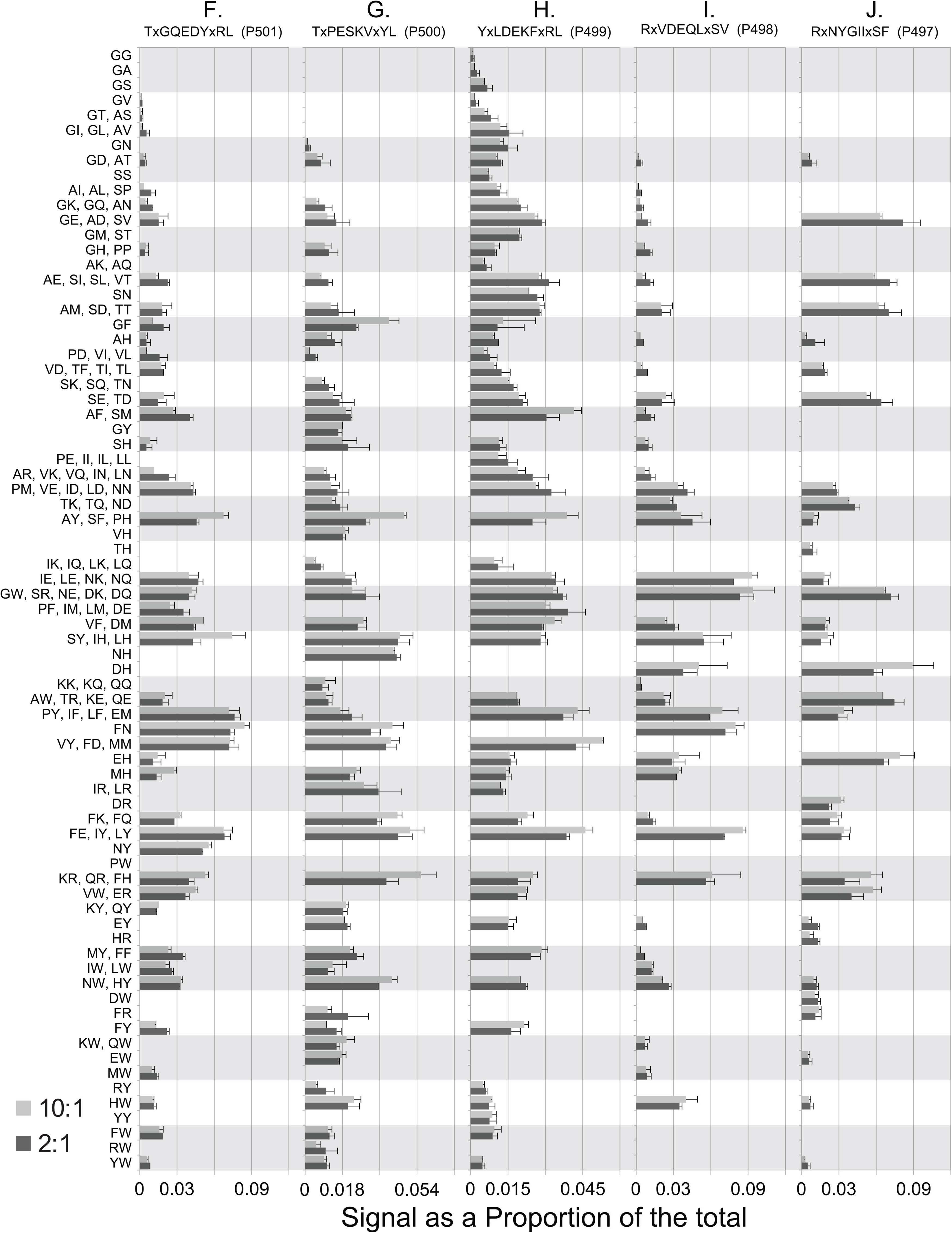
Panels F-J (continued from Fig. 9). Assembly assays with double-substitution peptide libraries based on five 11mer peptides (GHPDWYYGETW and four derivatives of the 11mer GHAEEYGAETL) and five 10mers (all from the original peptides found by gas phase sequencing, including the original REVDEQLLSV) show different preferences for particular amino acid combinations at P_2_, P_3_, P_c-3_ and P_c-2_ positions binding to BF2*21:01, with more combinations found for 10mer peptides. Bacterial-expressed heavy chain and β_2_-microglobulin were mixed with peptide libraries (19 amino acids, all except Cys, at one position, indicated by x in the peptide sequence) at two molar ratios (1:2:10 as usual, and 1:2:2 to control for strong binding peptides) and separated by SEC as described in the legend to Fig. 2 and separated immediately by SEC; monomer peaks were collected, concentrated, treated with acid, lyophilised and analysed by MALDI-TOF. Bar charts show proportion of total signal (with error bars when present showing standard error around the mean for separate refolding experiments) on x-axis for mass/charge (m/z) positions corresponding to peptides with particular amino acid combinations on the y-axis (single letter code; Ile and Leu, Gln and Lys, and amino acids present in P_2_ versus P_c-2_ positions were indistinguishable). Panels: F. TxGQEDYxRL, G. TxPESKVxYL, H. YxLDEKFxRL, I. RxVDEQLxSV, J. RxNYGIIxSF.

To get a more accurate understanding of the substitutions, six of these double-substitution libraries were analysed by LC-MS/MS, which allowed an exact identity and position of the amino acids in the combinations in peptides that refolded with BF2*21:01 (Fig. 11 – Source Data 1). The results by LC-MS/MS were amalgamated to give values that could be directly compared to the MALDI-TOF results (Figs. 11-13). Many combinations found by MALDI-TOF were also found by LC-MS/MS (ranging from 30-56.5% of total) with only a few found by MALDI-TOF but not by LC-MS/MS (ranging from 0-13% of total), while many more combinations were found by LC-MS/MS only (ranging from to 30.4-68.3% in groups that would have been equivalent to the MALDI-TOF identifications). The proportions of each combination by MALDI-TOF were higher than with LC-MS/MS, due to having fewer combinations in total. Thus, LC-MS/MS was more accurate and more sensitive than MALDI-TOF.

**Figure 11.**
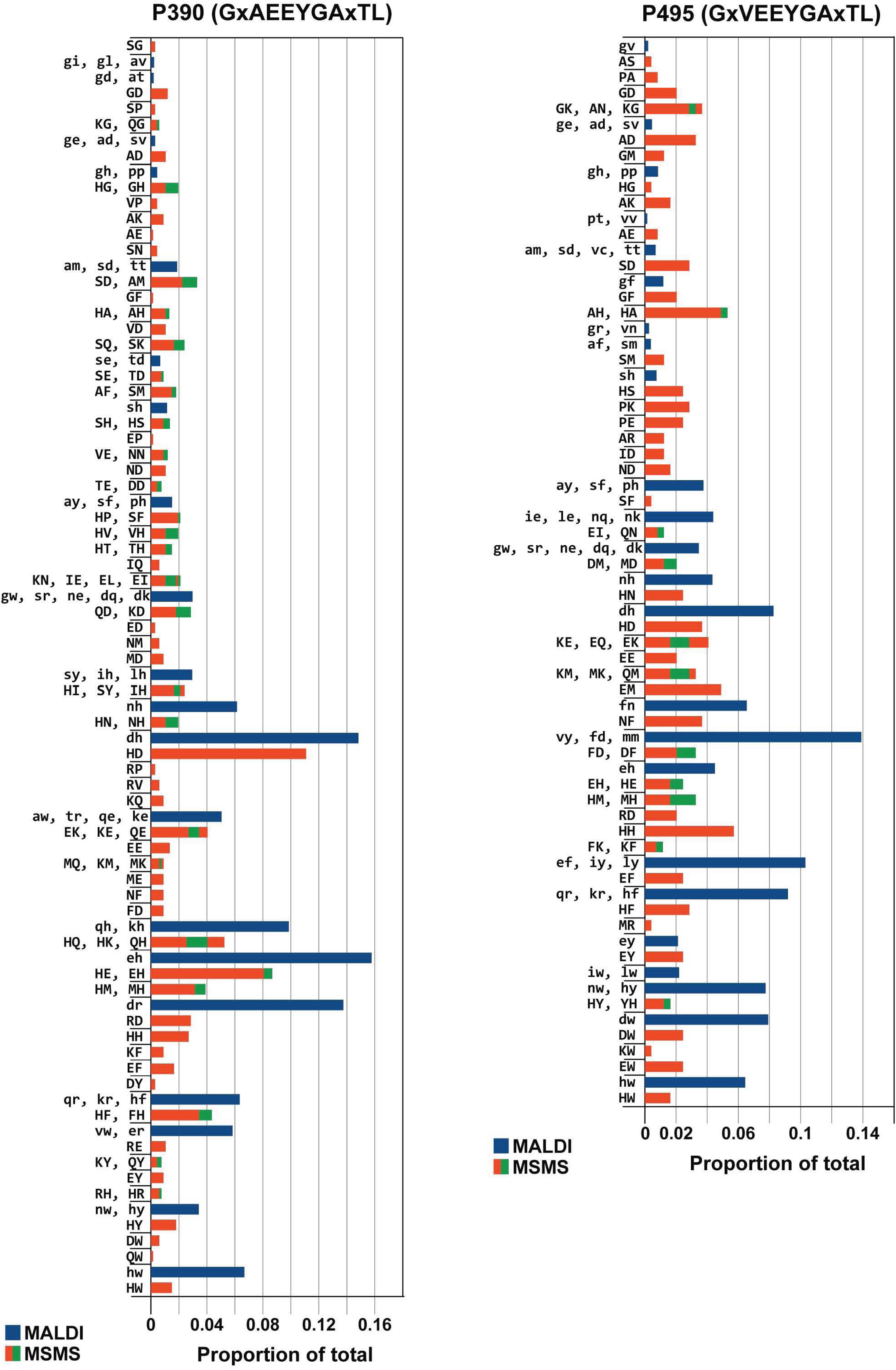
Panels A-B. Comparison of assembly assays of double-substitution libraries by analysis with MALDI-TOF and with LC-MS/MS give similar results, with different preferences for particular amino acid combinations at P_2_, P_3_, P_c-3_ and P_c-2_ positions binding to BF2*21:01, with more combinations for 10mer peptides. Bacterially-expressed heavy chain and β_2_-microglobulin were mixed with peptide libraries (19 amino acids, all except Cys, at two positions indicated by x in the peptide sequence) at two molar ratios (1:2:10 as usual, and 1:2:2 to control for strong binding peptides) and separated by SEC as described in the legend to Fig. 2; monomer peaks were collected, concentrated, treated with acid, lyophilised and analysed. Separate experiments were analysed by MALDI-TOF (blue bars, amino acids in single letter code with small letters, order not meaningful) and by LC-MS/MS (orange and green stacked bars, amino acids in single letter code with capital letters, first letter corresponding to P_2_ or P_3_ and second letter corresponding to P_c-2_ or P_c-3_, depending on peptide). Bar charts show proportion of total signal (with error bars when present showing standard error around the mean for separate refolding experiments) on x-axis for mass/charge (m/z) positions corresponding to peptides with particular amino acid combinations on the y-axis. Panels: A. GxAEEYGAxTL, B. GxVEEYGAxTL.

**Figure 12.**
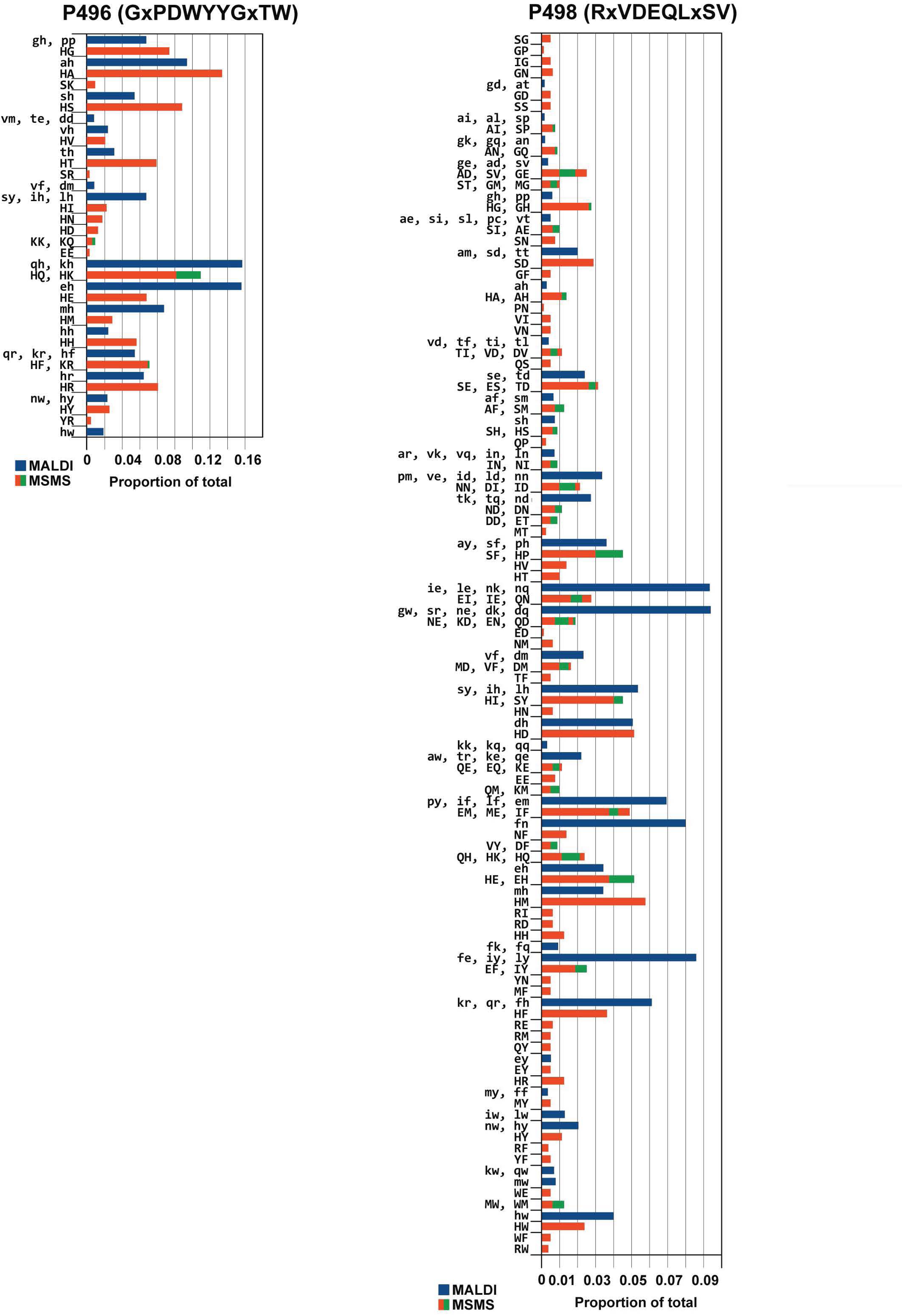
Panels C-D (continued from Fig. 11). Comparison of assembly assays of double-substitution libraries by analysis with MALDI-TOF and with LC-MS/MS give similar results, with different preferences for particular amino acid combinations at P_2_, P_3_, P_c-3_ and P_c-2_ positions binding to BF2*21:01, with more combinations for 10mer peptides. Bacterially-expressed heavy chain and β_2_-microglobulin were mixed with peptide libraries (19 amino acids, all except Cys, at two positions indicated by x in the peptide sequence) at two molar ratios (1:2:10 as usual, and 1:2:2 to control for strong binding peptides) and separated by SEC as described in the legend to Fig. 2; monomer peaks were collected, concentrated, treated with acid, lyophilised and analysed. Separate experiments were analysed by MALDI-TOF (blue bars, amino acids in single letter code with small letters, order not meaningful) and by LC-MS/MS (orange and green stacked bars, amino acids in single letter code with capital letters, first letter corresponding to P_2_ or P_3_ and second letter corresponding to P_c-2_ or P_c-3_, depending on peptide). Bar charts show proportion of total signal (with error bars when present showing standard error around the mean for separate refolding experiments) on x-axis for mass/charge (m/z) positions corresponding to peptides with particular amino acid combinations on the y-axis. Panels: C. GxPDWYYGxTW, D. RxVDEQLxSV.

**Figure 13.**
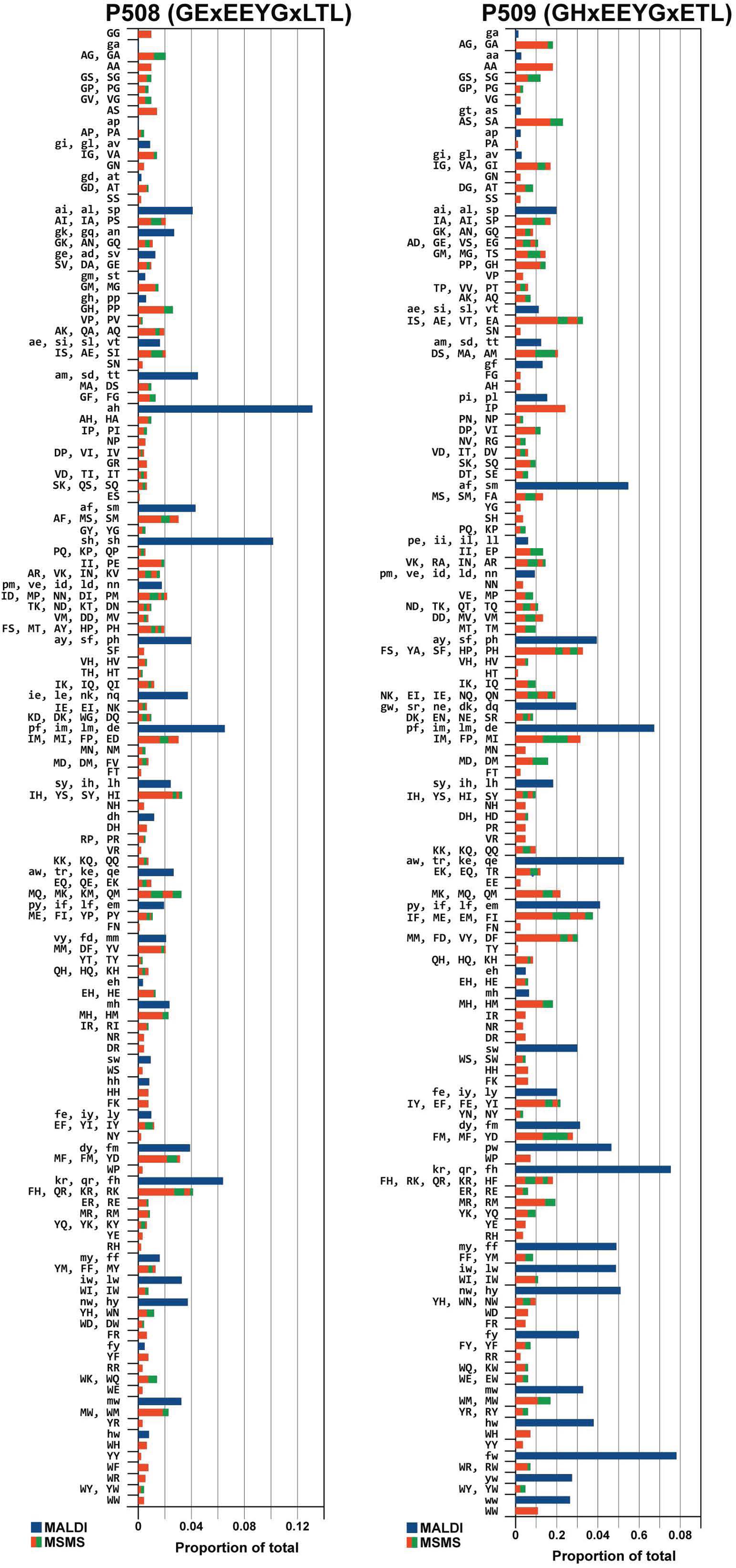
Panels E-F (continued from Figs. 11 and 12). Comparison of assembly assays of double-substitution libraries by analysis with MALDI-TOF and with LC-MS/MS give similar results, with different preferences for particular amino acid combinations at P_2_, P_3_, P_c-3_ and P_c-2_ positions binding to BF2*21:01, with more combinations for 10mer peptides. Bacterially-expressed heavy chain and β_2_-microglobulin were mixed with peptide libraries (19 amino acids, all except Cys, at two positions indicated by x in the peptide sequence) at two molar ratios (1:2:10 as usual, and 1:2:2 to control for strong binding peptides) and separated by SEC as described in the legend to Fig. 2; monomer peaks were collected, concentrated, treated with acid, lyophilised and analysed. Separate experiments were analysed by MALDI-TOF (blue bars, amino acids in single letter code with small letters, order not meaningful) and by LC-MS/MS (orange and green stacked bars, amino acids in single letter code with capital letters, first letter corresponding to P_2_ or P_3_ and second letter corresponding to P_c-2_ or P_c-3_, depending on peptide). Bar charts show proportion of total signal (with error bars when present showing standard error around the mean for separate refolding experiments) on x-axis for mass/charge (m/z) positions corresponding to peptides with particular amino acid combinations on the y-axis. Panels: E. GExEEYGxLTL, F. GHxEEYGxETL.

The LC-MS/MS results broadly follow those from the MALDI-TOF analysis (Figs. 11-13). Overall, there were 159 combinations at P_2_ and P_c-2_ found for double-substitution libraries based on four peptide backbones, with several (EE, HA, HD, HE, HF, HG, HH, HM, HN, HS and HY) found from all four libraries. However, there were more combinations found for the high affinity 10mer RxVDEQLxSV (75 combinations) than for the original 11mer GxAEEYGAxTL (60), the derivative GxVEEYGAxTL (44) and the low affinity 11mer GxPDWYYGxTW (23). The frequencies of amino acids in GxAEEYGAxTL and RxVDEQLxSV (Fig. 14) show that both prefer His at P_2_ (despite the original 10mer having Glu at P_2_) with a smattering of other amino acids including Glu and Ser. However, there were differences at P_c-2_, with the 11mer having Asp followed by Glu, His, Phe, Lys, Gln, Met and others at lower frequencies, but with the 10mer having nearly equal frequencies of Asp, Glu, Phe and Met, followed by Ile, Asn and others at lower frequencies. Moreover, the low affinity 11mer GxPDWYYGxTW has almost only His at P_2_, but a variety of amino acids at P_c-2_, most of which are poorly represented at P_c-2_ in GxAEEYGAxTL and RxVDEQLxSV (Fig. 15). Thus, the backbone of the peptide has effects on the preferences at P_c-2_.

**Figure 14.**
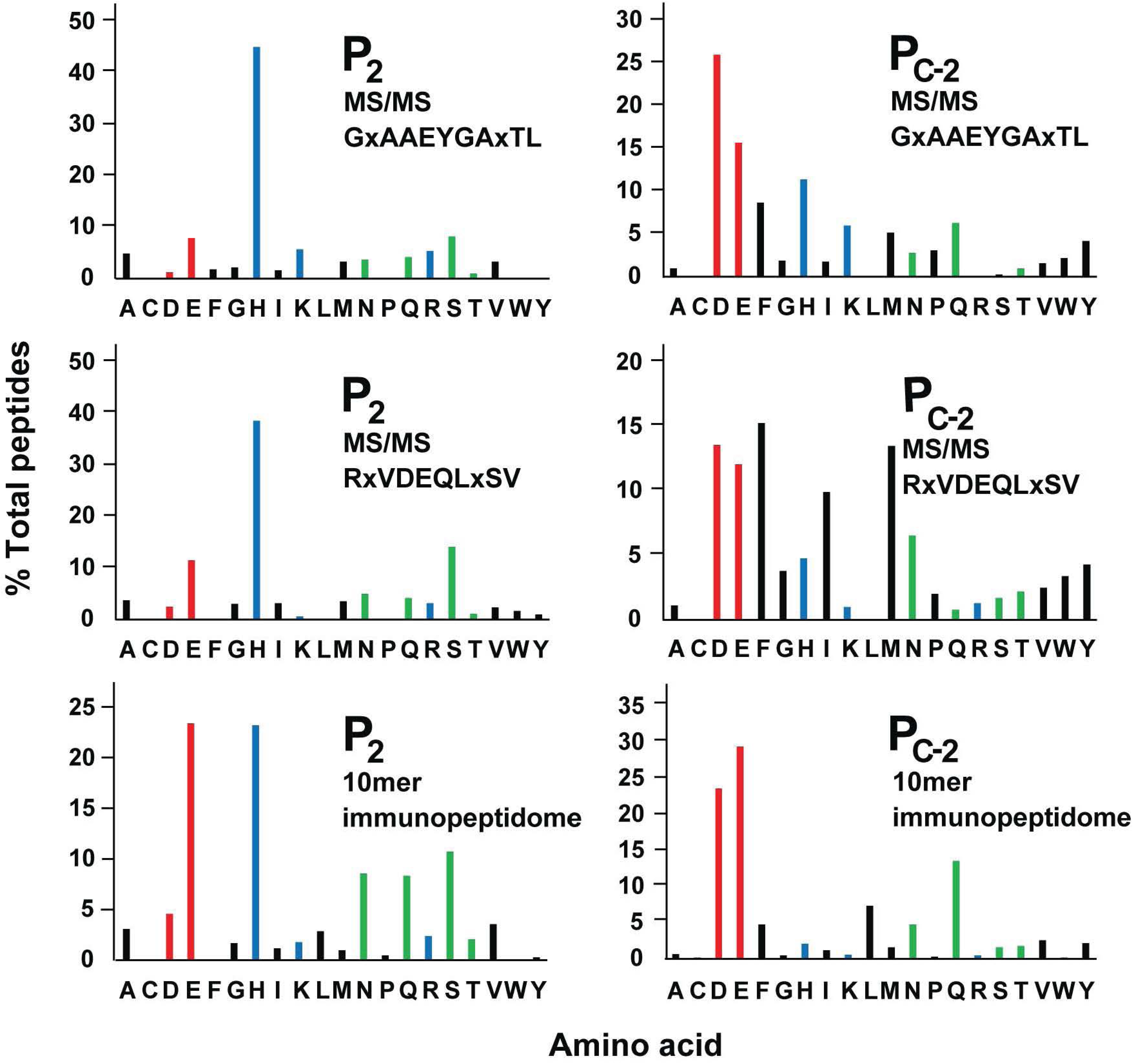
The frequencies of particular amino acids found at positions P_2_ and P_c-2_ of peptides bound to class I molecules from the B21 haplotype vary considerably, as assessed by LC-MS/MS of peptides from two double-substitution libraries with BF2*21:01 and from immunopeptidomics of a B21 cell line. For assembly assays, bacterial-expressed heavy chain and β_2_-microglobulin were refolded in vitro with either GxAAEYGAxTL or RsVDEQLxSV (single letter code; x means roughly equal proportions of 19 naturally-occurring amino acids, all except Cys), and separated by SEC; monomer peaks were collected, concentrated, treated with acid, and lyophilised. For immunopeptidomics, class I molecules from detergent-solubilised AVOL-1 cells were isolated by affinity chromatography with monoclonal antibody F21-2. Samples were analysed by LC-MS/MS, with bar graphs showing the percentage of sequences with different amino acids (single letter code) at P_2_ or P_c-2_ found by mass spectroscopy; immunopeptidomic data for 10mer peptides only.

**Figure 15.**
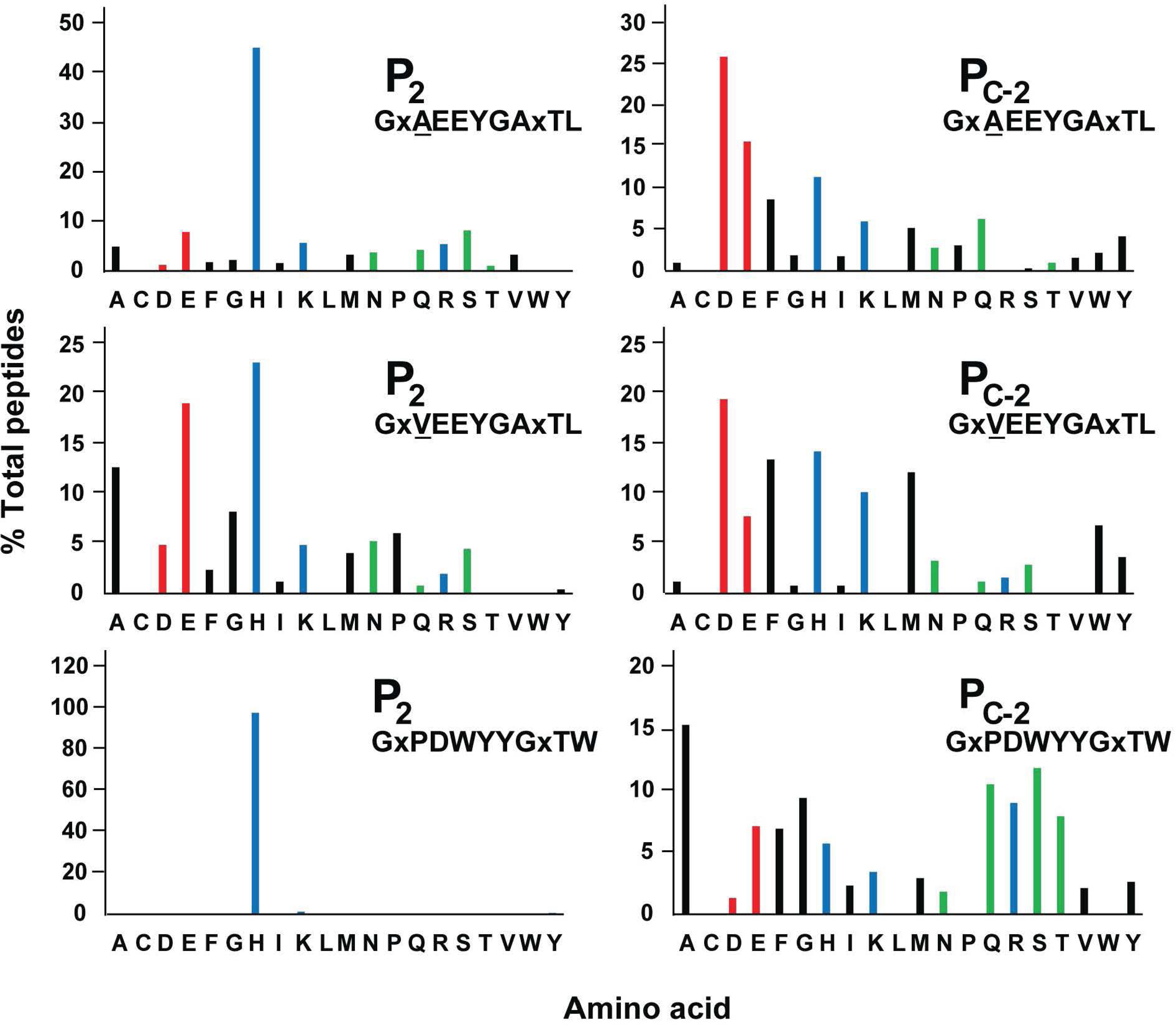
The frequencies of particular amino acids found at positions P_2_ and P_c-2_ in peptides bound to BF2*21:01 vary considerably and are affected by the amino acid present at P_3_, as assessed by LC-MS/MS of peptides from assembly assays with three double-substitution libraries. Bacterially-expressed heavy chain and β_2_-microglobulin were refolded in vitro with either GxAEEYGAxTL, GxVEEYGAxTL or GxPDWYYGxTW (single letter code; x means roughly equal proportions of 19 naturally-occurring amino acids, all except Cys) and separated by SEC as described in the legend to Fig. 2; monomer peaks were collected, concentrated, treated with acid, lyophilised and analysed by LC-MS/MS, with bar graphs showing the percentage of sequences with different amino acids (single letter code) at P_2_ or P_c-2_ found by mass spectroscopy.

The differences between GxAEEYGAxTL and GxVEEYGAxTL for LC-MS/MS and MALDI-TOF support the idea that P_3_ is an important position within the peptide (Figs. 8-13) despite not being an anchor reside (Chappell et al., 2015; Koch et al., 2008). In particular, the predominant His at P_2_ in the original 11mer with Ala at P_3_ changes to a more equal amount of His and Glu with significant Ala at P_2_ in the derivative with a Val at P_3_. Both LC-MS/MS and MALDI-TOF showed many combinations for P_3_ and P_c-3_ in the original 11mer GHxEEYGxETL and the derivative GExEEYGxLTL; the frequencies of amino acids at P_3_ were similar (with an increase Gly in the derivative), but several differences were found for P_c-3_ (including decreases in Lys, Phe, Arg and Ser in the derivative) (Fig. 16).

**Figure 16.**
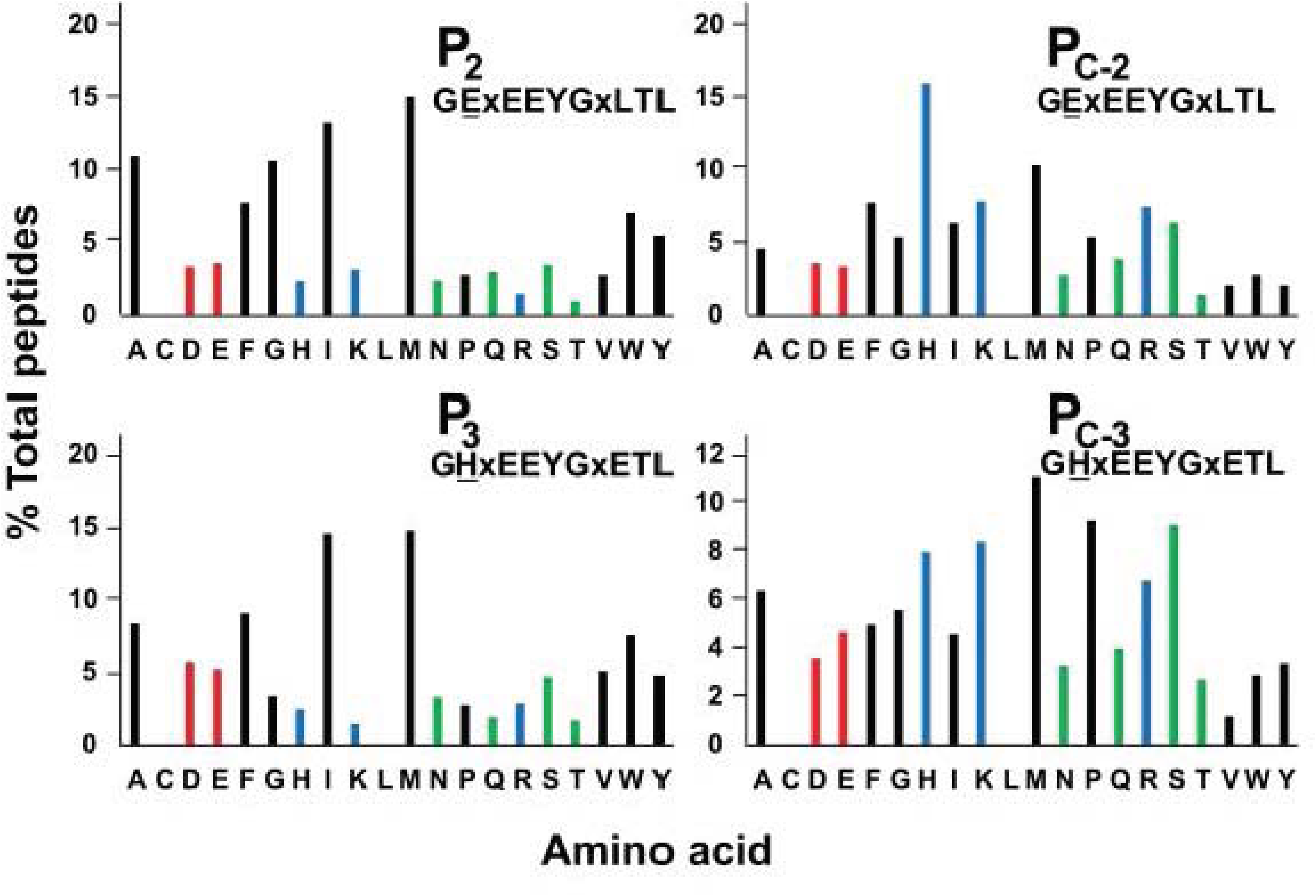
The frequency of particular amino acids found at positions P_3_ and P_c-3_ in peptides bound to BF2*21:01 is affected by the amino acids present at P_2_ and P_c-2_, as assessed by LC-MS/MS of peptides from assembly assays with two double-substitution libraries. Bacterially-expressed heavy chain and β_2_-microglobulin were refolded in vitro with either GExEEYGxLTL or GHxEEYGxETL (single letter code; x means roughly equal proportions of 19 naturally-occurring amino acids, all except Cys) and separated by SEC as described in the legend to Fig. 2; monomer peaks were collected, concentrated, treated with acid, lyophilised and analysed by LC-MS/MS, with bar graphs showing the percentage of sequences with different amino acids (single letter code) at P_3_ or P_c-3_ found by mass spectroscopy.

### Immunopeptidomics shows wide variation but only a few predominant combinations in anchor residues P_2_ and P_c-2_

In comparison with in vitro assembly assays, the peptides loaded within a cell are subject to quality control mechanisms, so the patterns found from in vitro and cellular analyses need not be the same. Immunopeptidomic analysis (Purcell et al., 2019) was used for the peptides eluted from class I molecules expressed in chicken cell lines, with a wide variety of peptides found for B21 compared to B19 (Chappell et al., 2015). There are two class I molecules in these preparations, but the expression of BF1 is much lower than BF2 (Shaw et al., 2008; Wallny et al., 2006). Immunopeptidomics of the B21 cells identified 3818 peptides with a range from 5mers to 50mers (Fig. 17 – Source Data 1), but the most frequent were found as 8, 9, 10, 11 and 12mers (2972 peptides, 77.8% of total). For each of these lengths, the peptide motifs determined by Gibbs clustering were complex (Fig. 17), as was found by in vitro assays. The 10mers were by far the most frequent (1470 peptides, 49.5% of 8-12mers) (Fig. 17), consonant with the wider variety of anchor residues and the greater thermostability noted above (Figs. 6-16). Peptides shorter than 8mers are conceivably proteolytic products, but at least some of the peptides longer than 12mers may have been bound to class I molecules, either by extra-long bulging in the middle of the peptide (Burrows et al., 2006; Liu et al., 2012) or by extending out of the groove at the C-terminus as has been found for other chicken class I molecules (Xiao et al., 2018).

**Figure 17.**
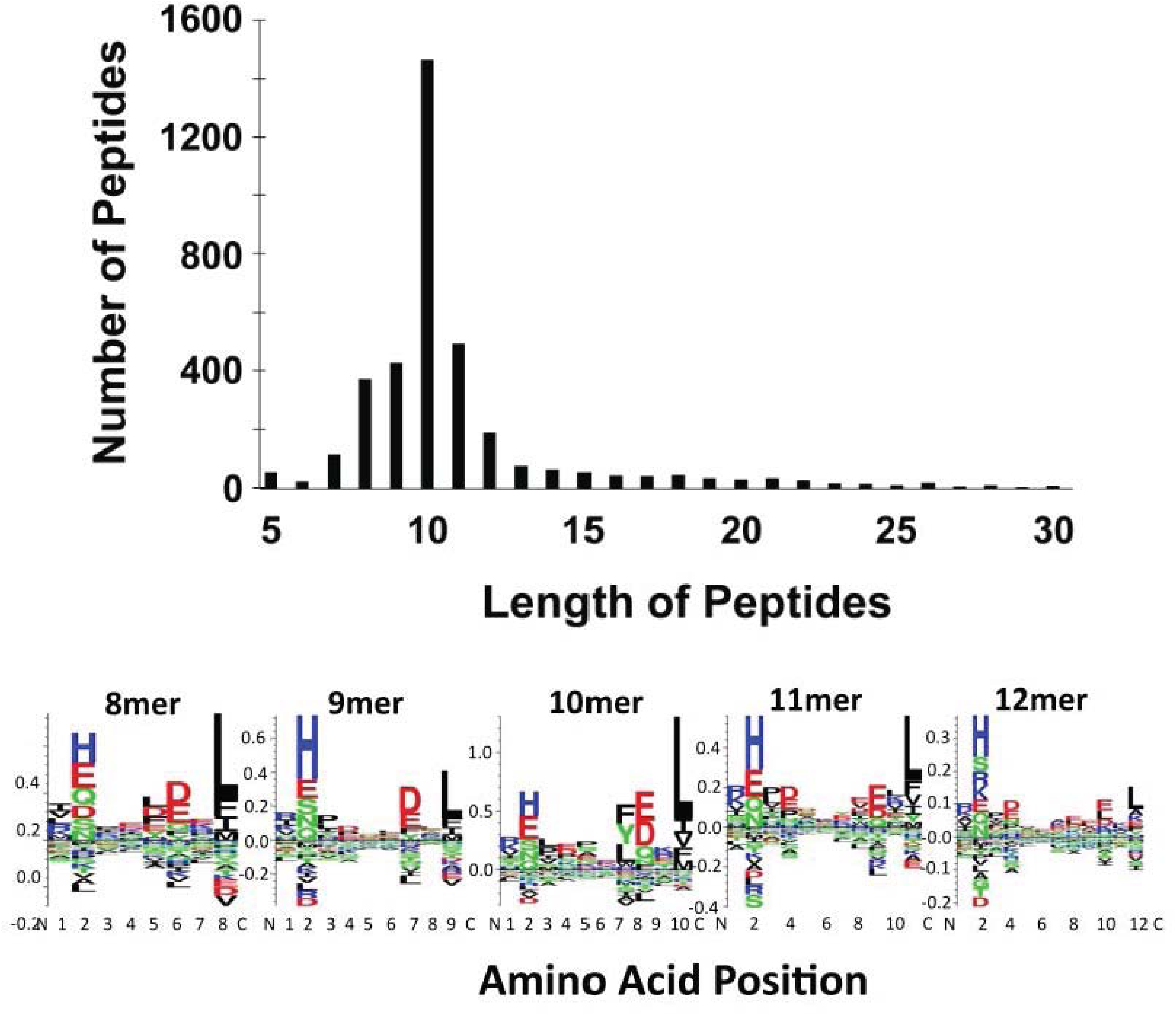
Most peptides found by immunopeptidomics of a B21 cell line were between 8 and 12 amino acids in length, with the most frequent being 10mers, all of which had complex peptide motifs. For immunopeptidomics, class I molecules from detergent-solubilised AVOL-1 cells were isolated by affinity chromatography with monoclonal antibody F21-2. Samples were analysed by LC-MS/MS, with bar graphs showing the number of sequences with different lengths as found by mass spectroscopy. Peptide motifs were determined by Gibbs clustering.

By immunopeptidomics, the frequencies of amino acids at key positions were similar but not identical for different peptide lengths (Fig. 18). For P_2_, the 10mers had roughly equal amounts of Glu and His followed by roughly equal levels of Asn, Gln and Ser and with other amino acids at lower levels. The 9mers and 11mers were enriched for His at P_2_ compared to 10mers, consistent with lower affinities of 9mers and 11mers not able to tolerate other amino acids so well. For P_c-2_, peptides of all three lengths prefer Asp and Glu, although they are roughly equal for 10mers, Asp favoured for 9mers and Glu favoured for 11mers; some other amino acids are more frequent in 10mers than 9mers or 11mers. The patterns for 12mers look like 9mers and 11mers, but for 8mers look more like 10mers (Fig. 18); studies in other haplotypes suggest that these 8mers may be at least in part from the BF1 molecule (data not shown). However, peptides of all five lengths overwhelmingly have Leu at P_c_ (Fig. 18).

**Figure 18.**
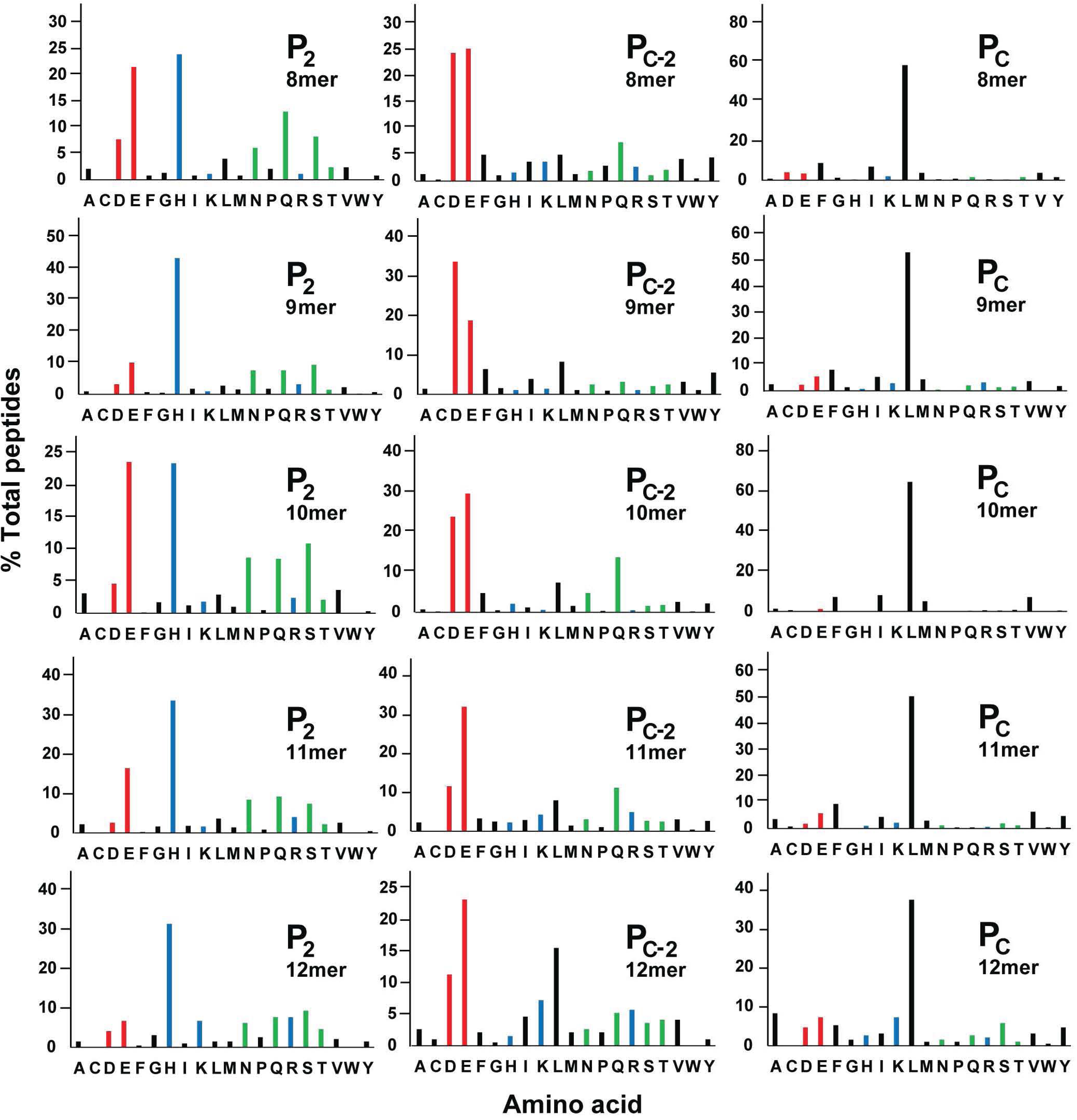
The frequency of particular amino acids found at positions P_2_, P_c-2_ and P_c_ in peptides bound to class I molecules from a B21 cell line varies depending on length of peptide, with Leu predominant at P_c_ for all lengths, and with 8mers more like 10mers at P_2_ and P_c-2_. Peptides identified by immunopeptidomics as in Fig. 17, with bar graphs showing the percentage of sequences with different amino acids (single letter code) at P_2_, P_c-2_ and P_c_ found by mass spectroscopy.

The analyses to resolve whether the peptides greater than 12mers bulged in the middle or hung out the C-terminal end gave equivocal results (Fig. 19). The 204 peptides from 13-15 amino acids in length had more His at P_2_ and Leu at P_c_ than other amino acids, but unlike the 12mers, the proportion of His at P_2_ was only marginally above many of the other amino acids and there was almost as much Lys as Leu at P_c_, with many other amino acids at significant amounts. The high levels of Asp, Glu and Leu at P_c-2_ found in 12mers might be expected to be located at P_c-2_ regardless of length if the peptides bulge in the middle, but might be expected at P_8_ regardless of length if a 10mer length is preferred and the rest of the peptide hangs out the end. Asp, Glu and Leu were high at both P_c-2_ and at P_8_, but many other amino acids were found there as well. It is possible that the several modes of binding contribute to the overall result, which were not possible to disentangle by these analyses.

**Figure 19.**
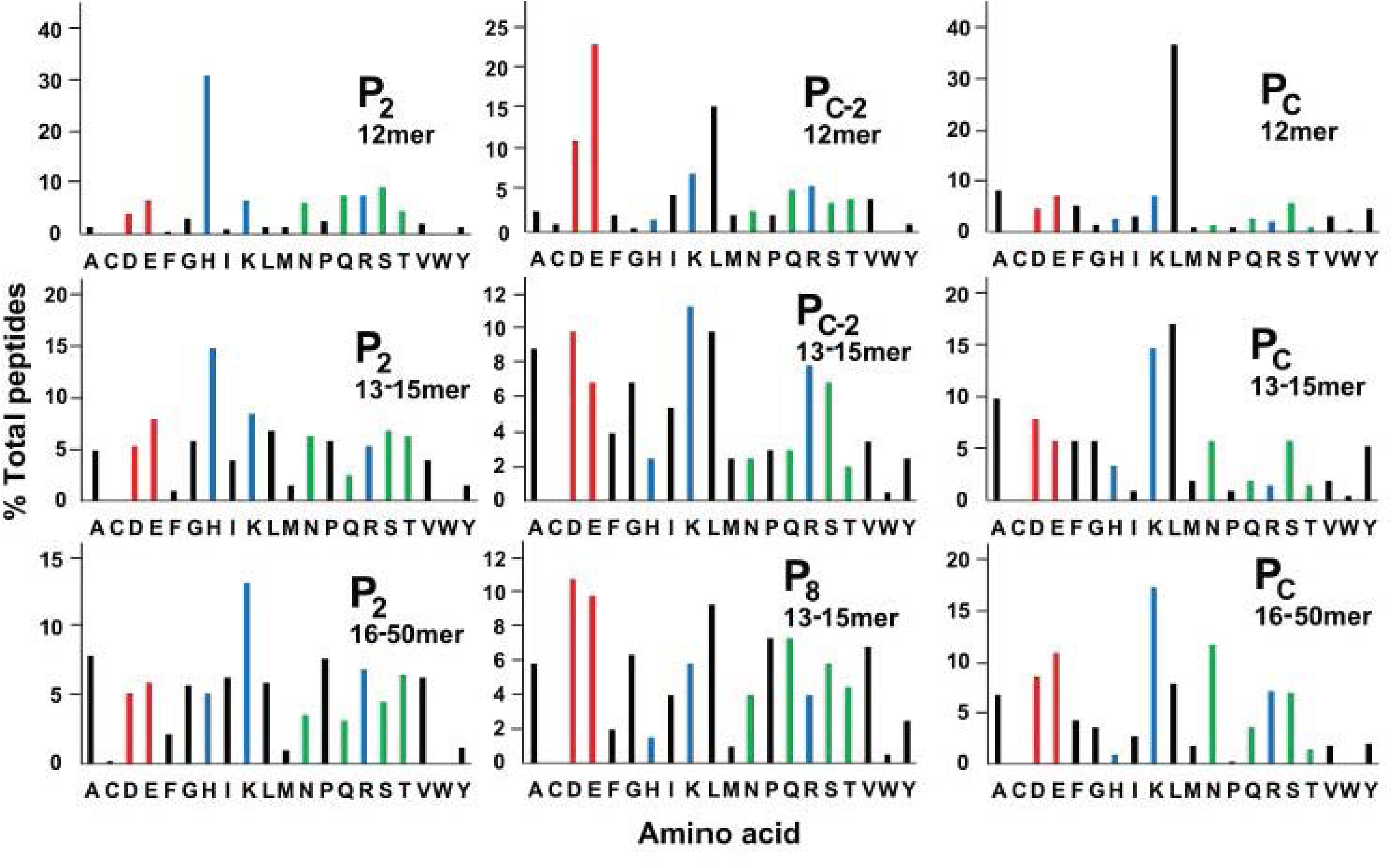
The frequency of particular amino acids found at positions P_2_, P_c-2_ and P_c_ in peptides bound to class I molecules from a B21 cell line varies depending on length of peptide, but patterns found in peptides up to 12mers are not so obvious in longer peptides. Peptides identified by immunopeptidomics as in Fig. 17, with bar graphs showing the percentage of sequences with different amino acids (single letter code) at P_2_, P_c-2_ and P_c_ found by mass spectroscopy.

Overall, the 442 peptides with lengths of 16 amino acids and above were even more different from the 12mers, with Lys favoured at both P_2_ and at P_c_ but with several other amino acids as well, and with the proportions of Asn, Glu and Ser at P_c_ increasing with increasing length.

The 10mer peptides had three groups of amino acids at P_2_ in roughly equal amounts: acidic (predominantly Glu), basic (predominantly His) or polar (roughly equally Asn, Gln and Ser with some Thr), with the remainder having hydrophobic amino acids at P_2_ (Fig. 20). The 10mers with acidic amino acids at P_2_ favoured Gln at P_c-2_ followed by Leu with small amounts of other amino acids, while the 10mers with basic, polar and hydrophobic amino acids at P_2_ favoured Asp and Glu at P_c-2_ with low levels of other amino acids, consonant with the co-variation at positions P_2_ and P_c-2_.

**Figure 20.**
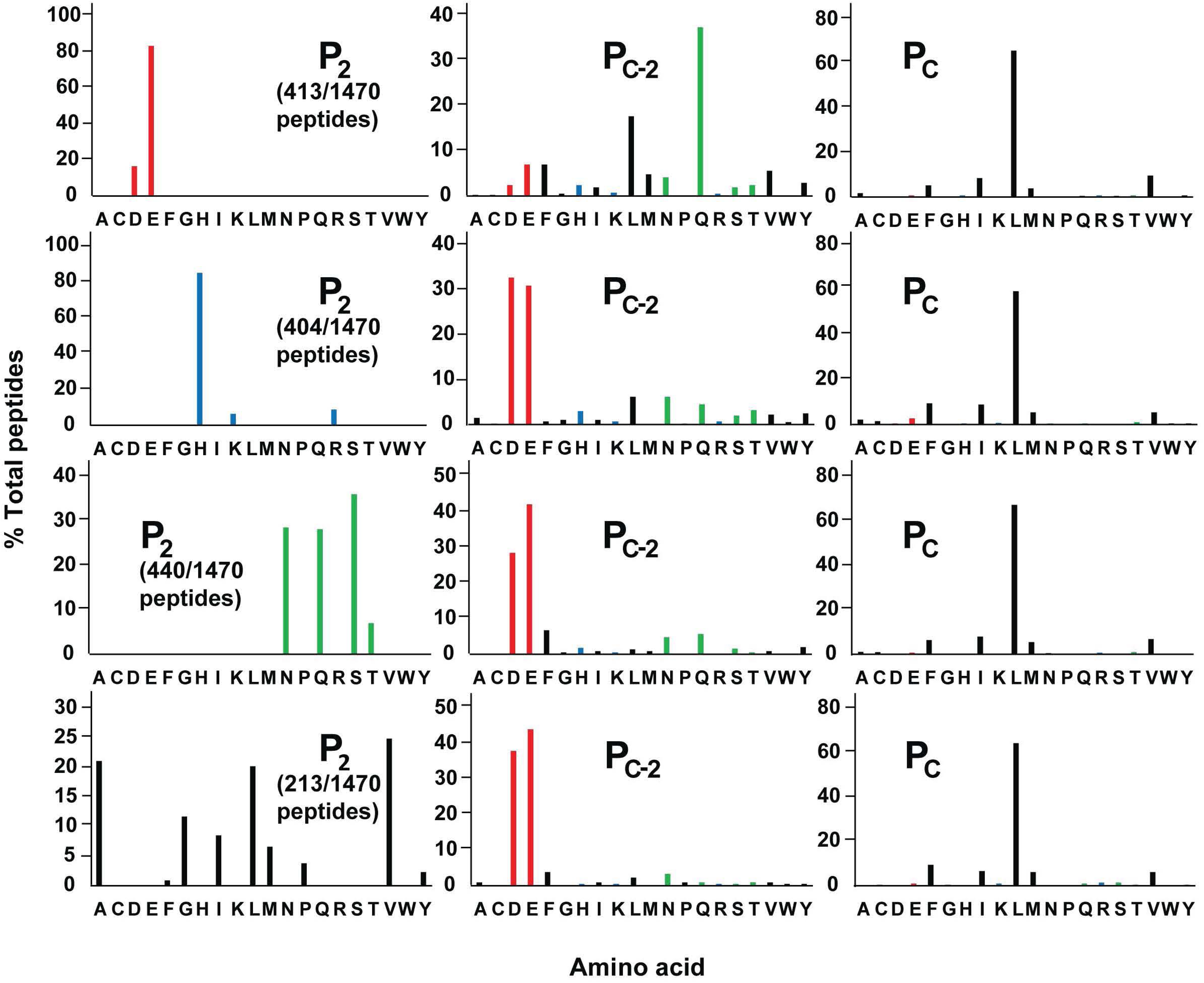
The frequencies of particular amino acids found at position P_c-2_ in peptides bound to class I molecules from a B21 cell line vary depending on whether the amino acids at P_2_, are acidic (Asp and Glu), basic (His, Lys and Arg), polar (Asn, Gln, Ser and Thr) or hydrophobic (Ala, Cys, Phe, Gly, Ile, Leu, Met, Pro, Val, Trp and Tyr), but Leu is found overwhelmingly at P_c_ in all peptides. Peptides identified by immunopeptidomics as in Fig. 17, with bar graphs showing the percentage of sequences with different amino acids (single letter code) at P_2_, P_c-2_ or P_c_ found by mass spectroscopy.

There were differences between the amino acid frequencies by immunopeptidomics and by assembly assays. In contrast to the predominant His at P_2_ for the assembly assays of both the 11mer GxAEEYGAxTL and 10mer RxVDEQLxSV, immunopeptidomics showed roughly equal Glu and His along with lesser amounts of Asn, Gln and Ser, followed by other amino acids (Fig. 14). In comparison to the complex patterns at P_c-2_ for the assembly assays, immunopeptidomics had predominant Asp and Glu with lesser amounts of Gln, Leu, Asn and Phe, followed by other amino acids (Fig. 14).

Co-variation between amino acids at P_2_ and P_c-2_ might be expected to reduce the number of combinations that bind to BF2*21:01, but 238 combinations were found for 9-11mers (273 for 8-12mers) out of 400 possible (Fig. 21). However, most of these combinations were found at very low levels, with the number of combinations found at greater than 1% frequency being only 19 for 9mers, 27 for 10mers and 17 for 11mers (Fig. 22); again 8mers were more like 10mers, perhaps due to the presence of peptides from BF1*2102. Of 36 high frequency combinations, 18 were shared among at least two of the 9mers, 10mers and 11mers (29 of 48 combinations if 8mers and 12mers are included). The combination His at P_2_ and Glu at P_c-2_ was predominant among 11mers at greater than 12% of total peptides, but His and Asp at nearly 16% was even more than His and Glu at 12% for the 9mers. A more even distribution was found for 10mers, with Glu and Gln at roughly 8%, His and Glu at over 7%, and His and Asp at nearly 7%; again 8mers were more like 10mers. Overall, those combinations above 1% frequency corresponded to 60% of 9mers, 71% of 10mers and 57% of 11mers, with the top two 9mers amounting to 27%, the top three 10mers to 22%, and the top 11mer to 12.4%. Thus, while a majority of combinations at P_2_ and P_c-2_ is possible, only a few were found frequently.

**Figure 21.**
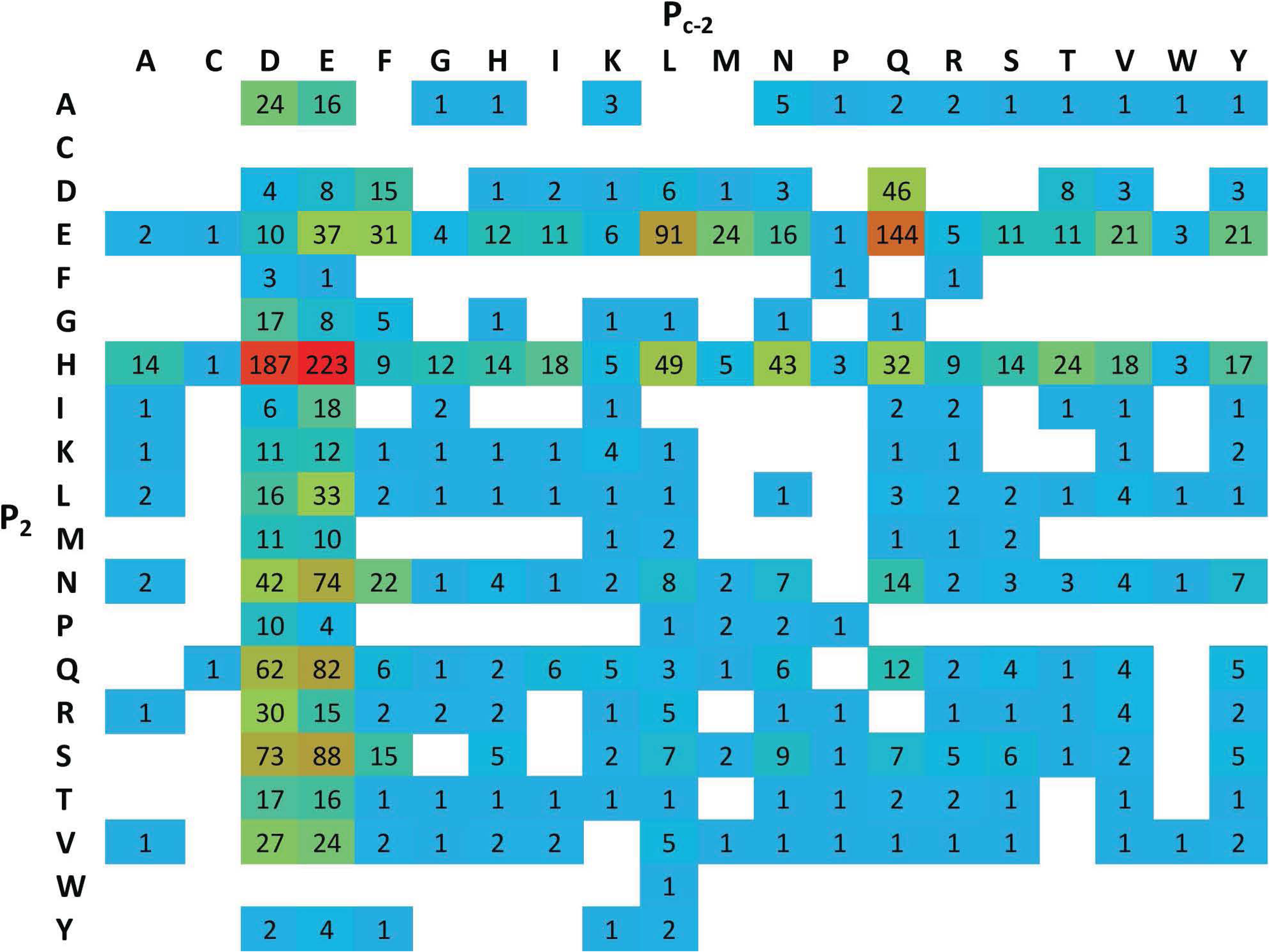
Many different combinations of amino acids at positions P_2_ and P_c-2_ in peptides bound to class I molecules from a B21 cell line are found by immunopeptidomics, but only a few are found frequently. Peptides (9, 10 and 11mers) identified by immunopeptidomics as in Fig. 17, with heat map showing combinations of amino acids (single letter code, with P_2_ on left and P_c-2_ on top) at intersections, with empty cells showing no peptides found, and colours showing numbers of peptides found highlighted with low frequency in blue to high frequency in red.

**Figure 22.**
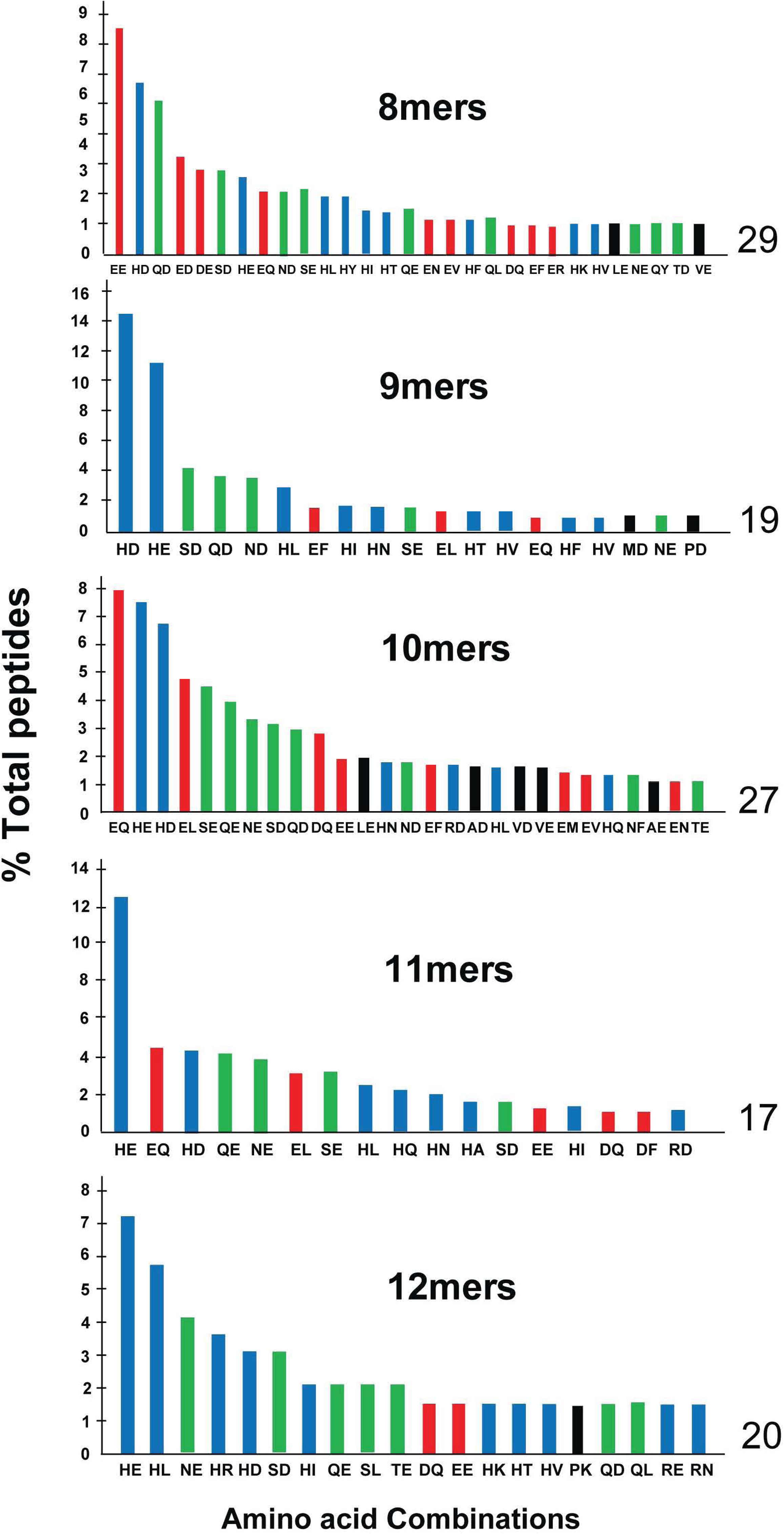
Only a few combinations of amino acids at positions P_2_ and P_c-2_ in peptides bound to class I molecules from a B21 cell line are found at frequencies of 1% or higher. Peptides identified by immunopeptidomics as in Fig. 17, with bar graphs showing the percentage of sequences with different combinations of amino acids at P_2_ and P_c-2_ (single letter code, with P_2_ first and then P_c-2_) found by mass spectroscopy.

### Immunopeptidomics shows wide variation and complex patterns for P_3_ and P_c-3_ that are affected by P_2_ and P_c-2_

Analysis from the immunopeptidomic data supports the interpretation from the assembly assays that the amino acids allowed at the anchor residues (in the sense of binding into deep pockets of the MHC molecule) at P_2_ and P_c-2_ may be affected by amino acid sidechains from positions that are not anchor residues. In particular, the amino acids at P_3_ and P_c-3_, which do not contact the MHC molecule at all (Chappell et al., 2015; Koch et al., 2008), are found at wildly varying proportions depending on peptide length (Fig. 23). Pro at P_3_ was found at high frequency (24% for 9mers, 17% for 10mers, 18% for 11mers) as well as Leu (10% for 9mers, 19% for 10mers, 13% for 11mers), with significant amounts of Ala, Ile, Met, Ser and Val for 10mers (and less for 9mers). At P_c-3_, Phe, Leu and Tyr were frequent (each at about 20%) with a smattering of other residues in 10mers, while in addition Ala, Asp, Glu Ile, Lys, Pro and Val were found half as often in the 9mer, and Glu was twice as likely as Ala, Asp, Phe, His, Ile, Leu, Ser, Val and Trp in the 11mers. The same amino acids were found for 8mers but at different proportions (Fig. 23), which may indicate binding to BF1 rather than BF2.

**Figure 23.**
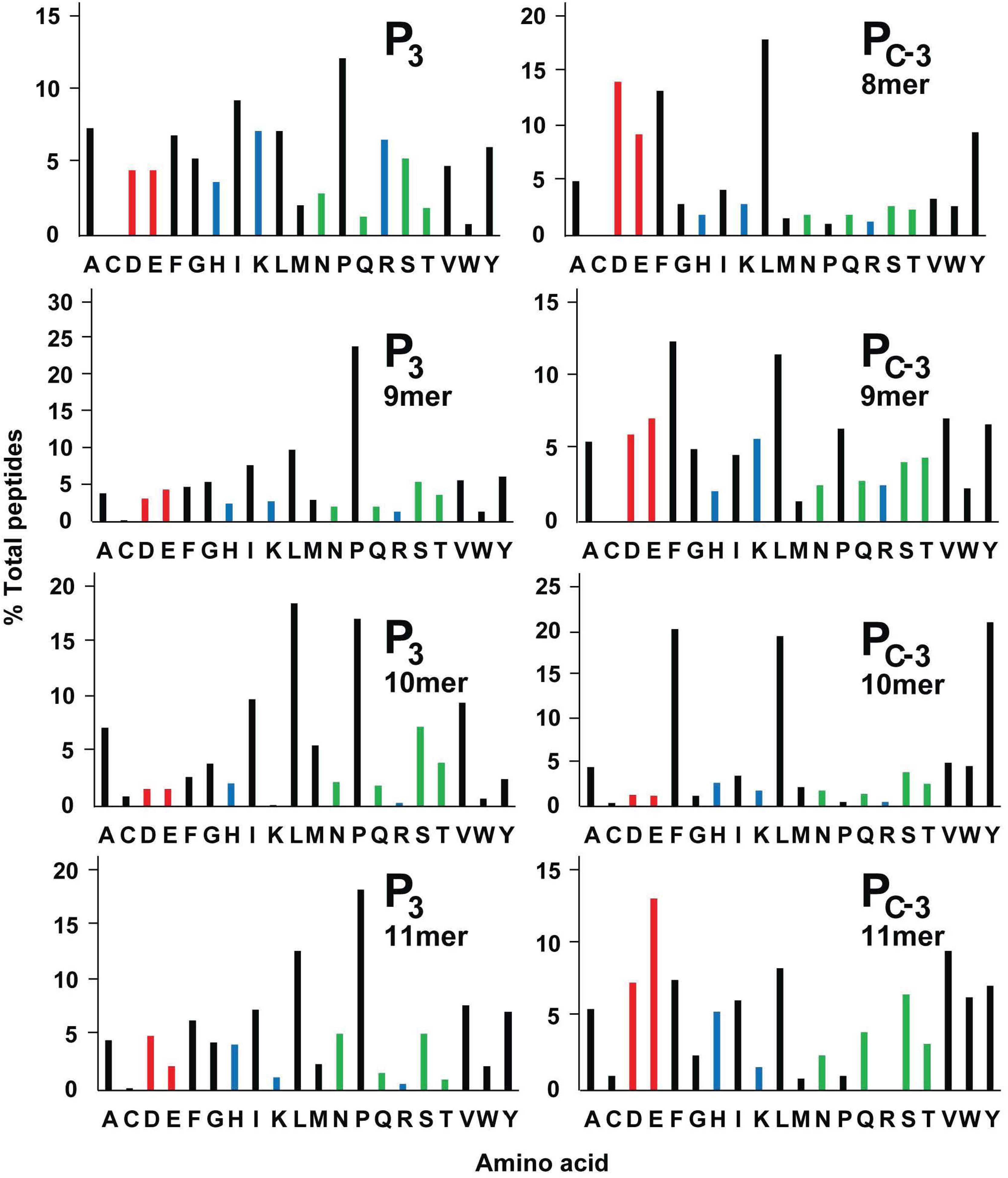
The frequencies of particular amino acids found at positions P_3_ and P_c-3_ in peptides bound to class I molecules from a B21 cell line vary depending on length of peptide. Peptides identified by immunopeptidomics as in Fig. 17, with bar graphs showing the percentage of sequences with different amino acids (single letter code) at P_3_ or P_c-3_ found by mass spectroscopy.

A more detailed look at the immunopeptidomic data suggests that the amino acids allowed at positions P_3_ and P_c-3_ may depend on the amino acids at P_2_ and P_c-2_ (Figs. 24, 25). For example, acidic residues (Asp and Glu) at P_3_ are found at high levels only in 11mers with Tyr at P_2_, in 10mers with Phe (and less so for Tyr) at P_2_, and in 9mers with several amino acids at P_2_ (Fig. 24). In contrast, basic residues at P_3_ are nearly all found in 11mers with Phe at P_2_, in over 80% of 9mers with Ile, Leu and Tyr at P_2_, but much more evenly distributed in 10mers. Large hydrophobic residues (Phe, Ile, Leu, Met and Tyr) at P_3_ are found at high levels in many peptides, but not 9mers with Ala, Phe, Gly, Ile, Lys and Met at P_2_, in 10mers with Phe and Pro at P_2_, and in 11mers with Ala, Phe and Tyr at P_2_ (as well as Cys and Trp, but there are not many peptides total for these residues). Similarly, there are few if any peptides with small hydrophobic residues (Ala, Cys, Gly and Ser) at P_3_ for 9mers with Asp, Glu and Tyr, and for 11mers with Phe, Lys, Pro and Tyr. The kinked amino acid Pro at P_2_ is found in nearly 70% of 10mers with Pro at P_3_, and strong (but not so extreme) skewing is found for 9mers and 11mers. For example, for 9mers with Pro at P_3_ there at high levels of Phe, Gly, His, Met, Asn, Ser and Thr at P_2_, but none with Lys, Leu, Arg or Tyr. Such unequal distributions of P_c-3_ dependent on P_c-2_ are found for 9mers, 10mer and 11mers as well (Fig. 25). Particularly striking are the extreme preferences in 9mers for Arg at P_c-2_ with acidic residues at P_c-3_, Pro at P_c-2_ with basic resides at P_c-3_, and Ala at P_c-2_ with Pro at P_c-3_. Thus, whether a particular peptide binds BF2*21:01 depends not only on the anchor residues at P_2_, P_c-2_ and P_c_, but also on residues from the rest of the peptide, such as P_3_ and P_c-3_.

**Figure 24.**
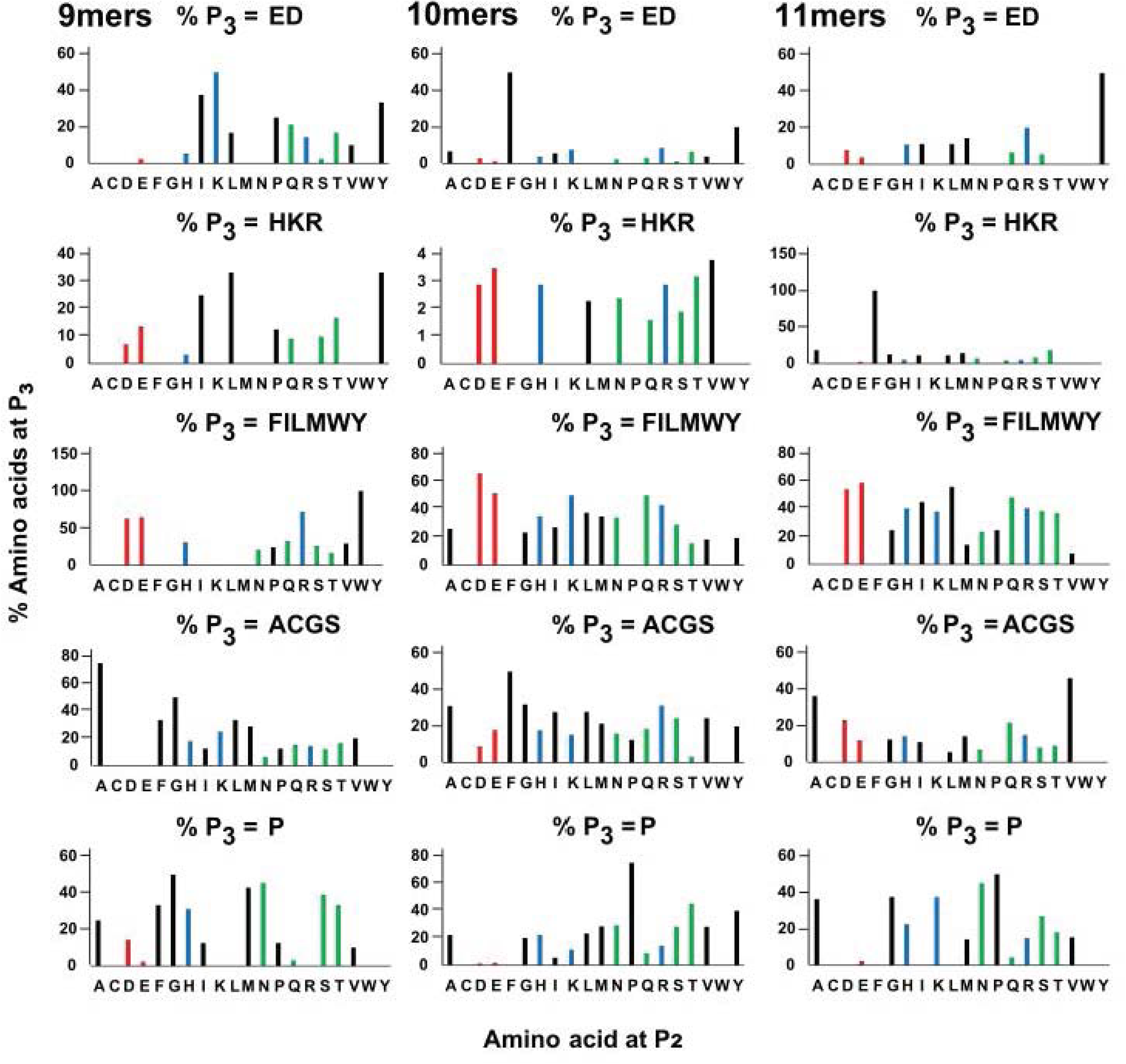
The frequencies of groups of amino acids (acidic, basic, large hydrophobic, small and Pro) found at position P_3_ in peptides bound to class I molecules from a B21 cell line vary depending on the amino acid at P_2_ and on the length of peptide. Peptides identified by immunopeptidomics as in Fig. 17, with bar graphs showing the percentage of sequences with amino acids at P_2_ on the x-axis against different groups of amino acids (single letter code: acidic being Asp and Gly; basic being His, Lys and Arg; large hydrophobic being Phe, Ile, Leu, Met, Trp and Tyr; small being Ala, Cys, Gly Ser; Pro) at P_3_ on the y-axis, as found by mass spectroscopy.

**Figure 25.**
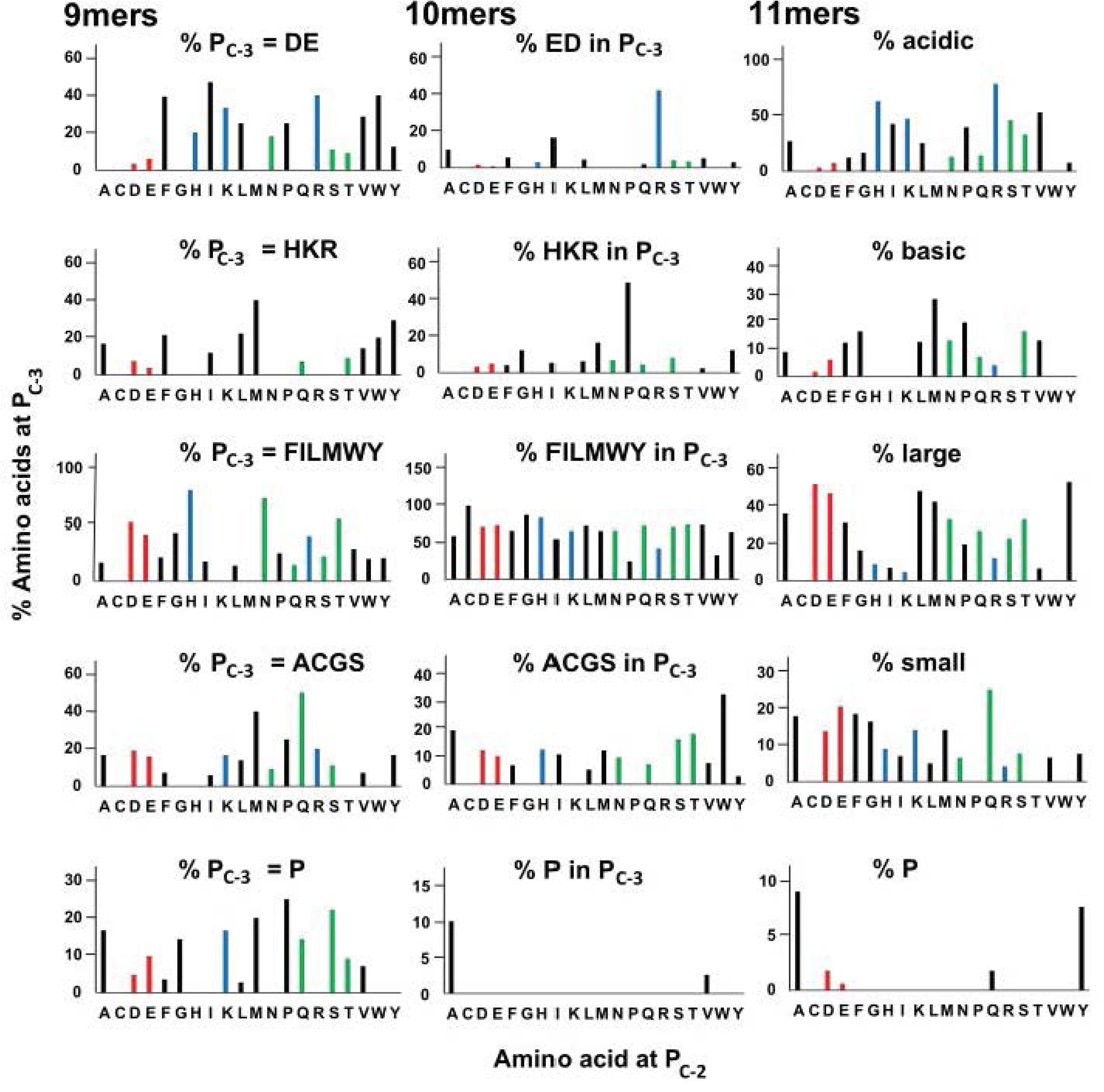
The frequencies of groups of amino acids (acidic, basic, large hydrophobic, small and Pro) found at positions P_c-3_ in peptides bound to class I molecules from a B21 cell line vary depending on the amino acid at P_c-2_ and on length of peptide, with 8mers more like 10mers. Peptides identified by immunopeptidomics as in Fig. 17, with bar graphs showing the percentage of sequences with amino acids at P_c-2_ on the x-axis against different groups of amino acids (single letter code: acidic being Asp and Gly; basic being His, Lys and Arg; large hydrophobic being Phe, Ile, Leu, Met, Trp and Tyr; small being Ala, Cys, Gly Ser; Pro) at P_c-3_ on the y-axis, as found by mass spectroscopy.

### Immunopeptidomics shows a strong preference for Leu at P_c_ but co-variation with P_c-2_

Assembly assays with GHAEEYGAETx and REVDEQLLSx (Fig. 7) show high and roughly equal amounts of Phe, Ile/Leu, Met and Val at P_c_, but immunopeptidomics shows Leu at P_c_ to be by far the most frequent, 50-60% frequency in 8-11mers (Fig. 18). At much lower levels are found Ile, Phe, Met and Val, but in fact every single other amino acid is found at P_c_ by immunopeptidomics. The reason underlying this strong preference for Leu at Pc in a living cell compared to assembly assays is not clear, but one possibility is co-variation with other peptide positions (unlike the assembly assays with fixed sequences).

Peptides with different amino acids at P_c-2_ can have much different preferences for amino acids at P_c_ (Figs. 26, 27). The levels of Leu at P_c_ are greater than 50% for five amino acids in 9mers, 12 for 10mers and six for 11mers, with Asp, Glu and Gln at P_c-2_ having high Leu frequencies at P_c_ for all three lengths. In contrast, the percentages of Leu at P_c_ are low for Arg at P_c-2_ in 9mers, Arg and Lys for 10mers, and Arg, Lys and Ile for 11mers (Fig. 26).

**Figure 26.**
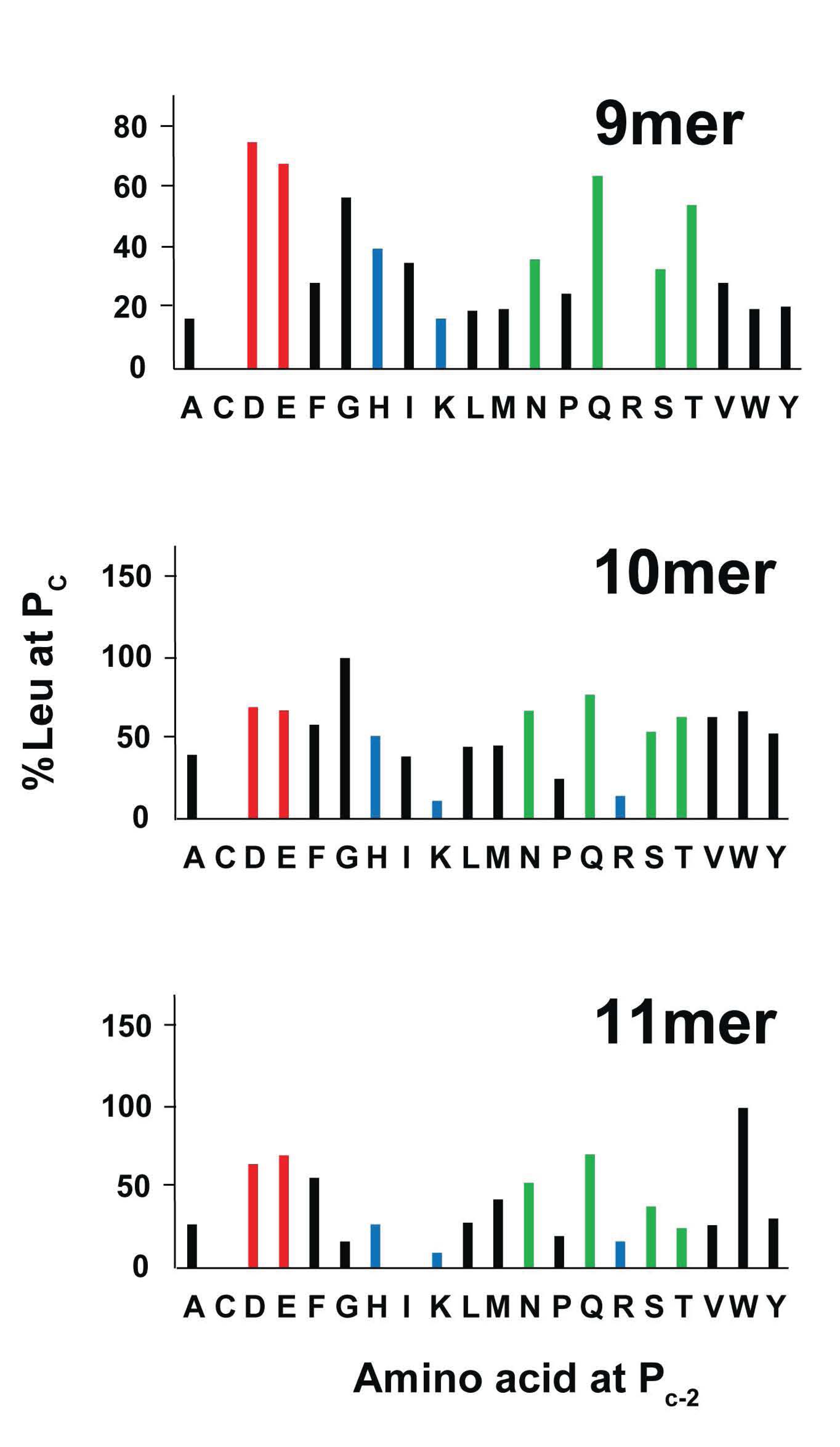
The frequency of Leu found at position P_c_ in peptides bound to class I molecules from a B21 cell line vary depending on length of peptide and position P_c-2_. Peptides identified by immunopeptidomics as in Fig. 17, with bar graphs showing the percentage of sequences with Leu at P_c-2_ on the y-axis against amino acids (single letter code) at P_c_ on the y-axis, as found by mass spectroscopy.

For many amino acids at P_c-2_, there is a preponderance of Leu at P_c_ followed by other hydrophobic amino acids at P_c_. For 10mers, the presence of Leu and other hydrophobic amino acids is almost complete. In contrast, 9mers with Ala, Phe, Leu, Met, Pro, Ser, Trp and Tyr as well as 11mers with Gly, His, Ile, Lys and Pro at P_c_ have greater than 35% acidic (Asp or Glu), basic (Arg, His or Lys) or polar (Asn, Gln, Ser or Thr) amino acids at P_c-2_ (Fig. 27). For example, out of the 28 9mer sequences with Phe at P_c-2_, only eight have Leu at P_c_, but 13 (46.4%) have acidic, basic or polar amino acids. This compares to 70 10mer sequences with Phe at P_c-2_, of which 41 have Leu at P_c_ and only five are acidic, basic or polar. Similarly, out of the 24 9mer sequences with Tyr at P_c-2_, only five have Leu at P_c_, but 15 (62.5%) have acidic, basic or polar amino acids; of 32 such sequences that are 10mers, 17 have Leu at P_c_ and only seven are acidic, basic or polar. Although there are only four 9mers, four 10mers and five 11mers with Pro at P_c-2_, only one sequence each has Leu at P_c_, while 50-75% are acidic, basic or polar. For sequences with Leu at P_c-2_, 14 out of 36 9mers (39%), 12 out of 39 11mers (36%), and even 13 out of 109 10mers (12%) have acidic, basic or polar amino acids at P_c_.

**Figure 27.**
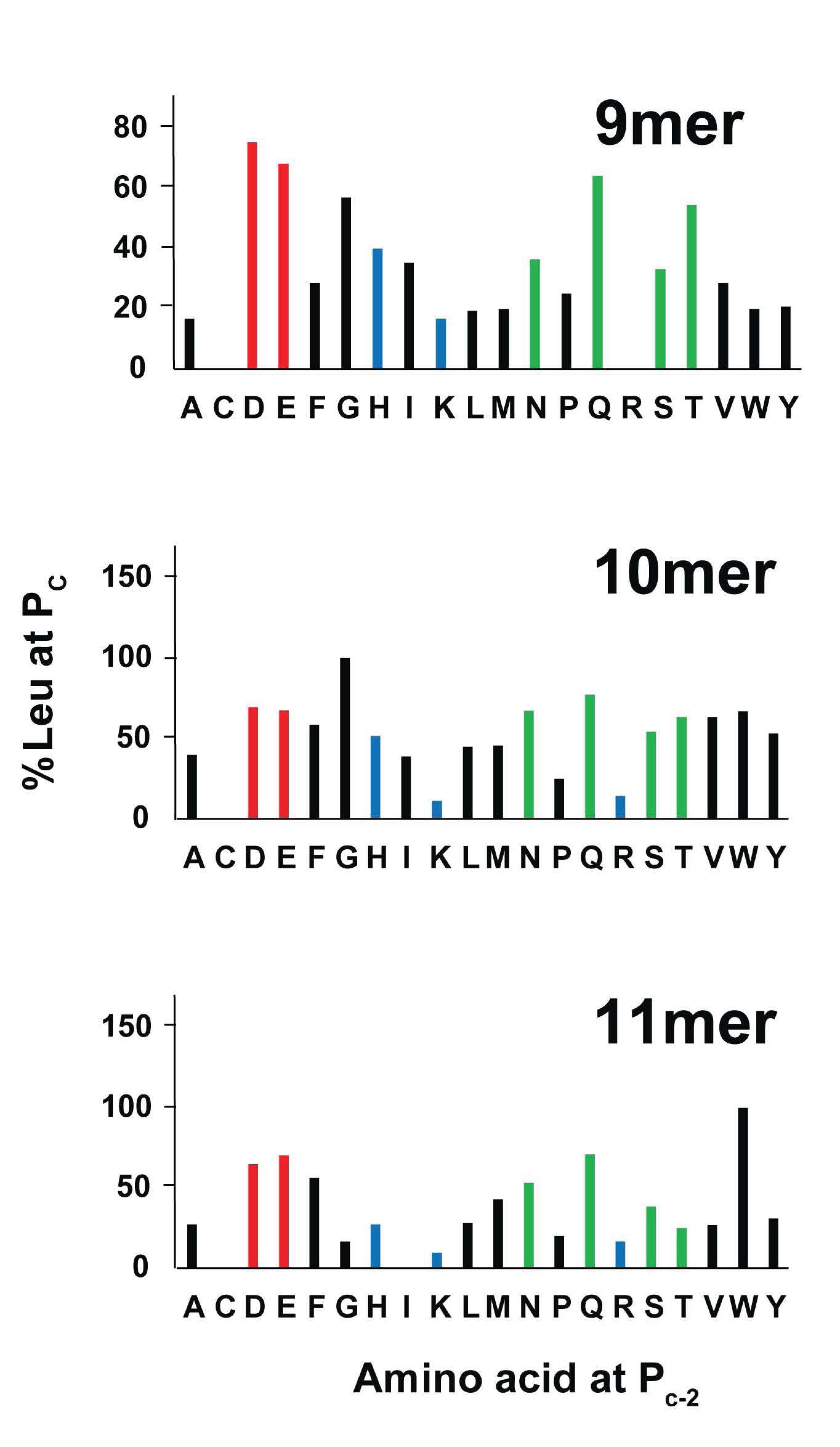
The frequencies of groups of amino acids (Leu, other hydrophobic, acidic, basic, polar) found at P_c_ in peptides bound to class I molecules from a B21 cell line vary depending on the amino acid at P_c-2_ and on length of peptide, with significant proportions of acidic, basic and/or polar amino acids compared to Leu at Pc for certain amino acids at P_c-2_ in 9mers and 11mers. Peptides identified by immunopeptidomics as in Fig. 17, with stacked bar graphs showing the number of sequences with Leu, other hydrophobic (Cys, Phe, Gly, Ile, Met, Pro, Val, Trp and Tyr), acidic (Asp and Glu), basic (His, Lys and Arg) and polar (Asn, Gln, Ser and Thr) amino acids at P_c_ against length of peptide and amino acid (single letter code) at P_c-2_, as found by mass spectroscopy.

Such associations suggest that co-variation also occurs between the amino acids at P_c-2_ and P_c_, presumably through the interactions with BF2*21:01 rather than directly. It is also clear (Fig. 24) that the high frequency of 10mers and of Asp and Glu at P_c-2_ among peptides found on BF2*21:01 molecules favours overall the presence of Leu at P_c_.

### The C-terminal part of peptides bound to BF2*21:01 have very similar conformations

Thus far, eight structures have been determined for 10mer and 11mer peptides bound to the dominantly-expressed class I molecule of the B21 haplotype, BF2*21:01. Despite the very wide range of residues found for the anchor residues P_2_ and P_c-2_, as well as the non-anchor residues P_3_ and P_c-3_, the main chains of these eight peptides are nearly superimposable for the seven amino acids at the C-terminus (P_c-4_ through P_c_), including the descending helical region of P_c-5_ through P_c-3_ (Fig. 28). In contrast, the conformations in the N-terminal end of the peptide (P_2_ and P_3_ for 10mers, P_2_ through P_4_ for 11mers, and perhaps including P_1_) vary significantly. B factors are also higher for the main chain atoms of the N-terminal part of the peptide compared to the C-terminal part, signifying greater flexibility in the structures.

**Figure 28.**
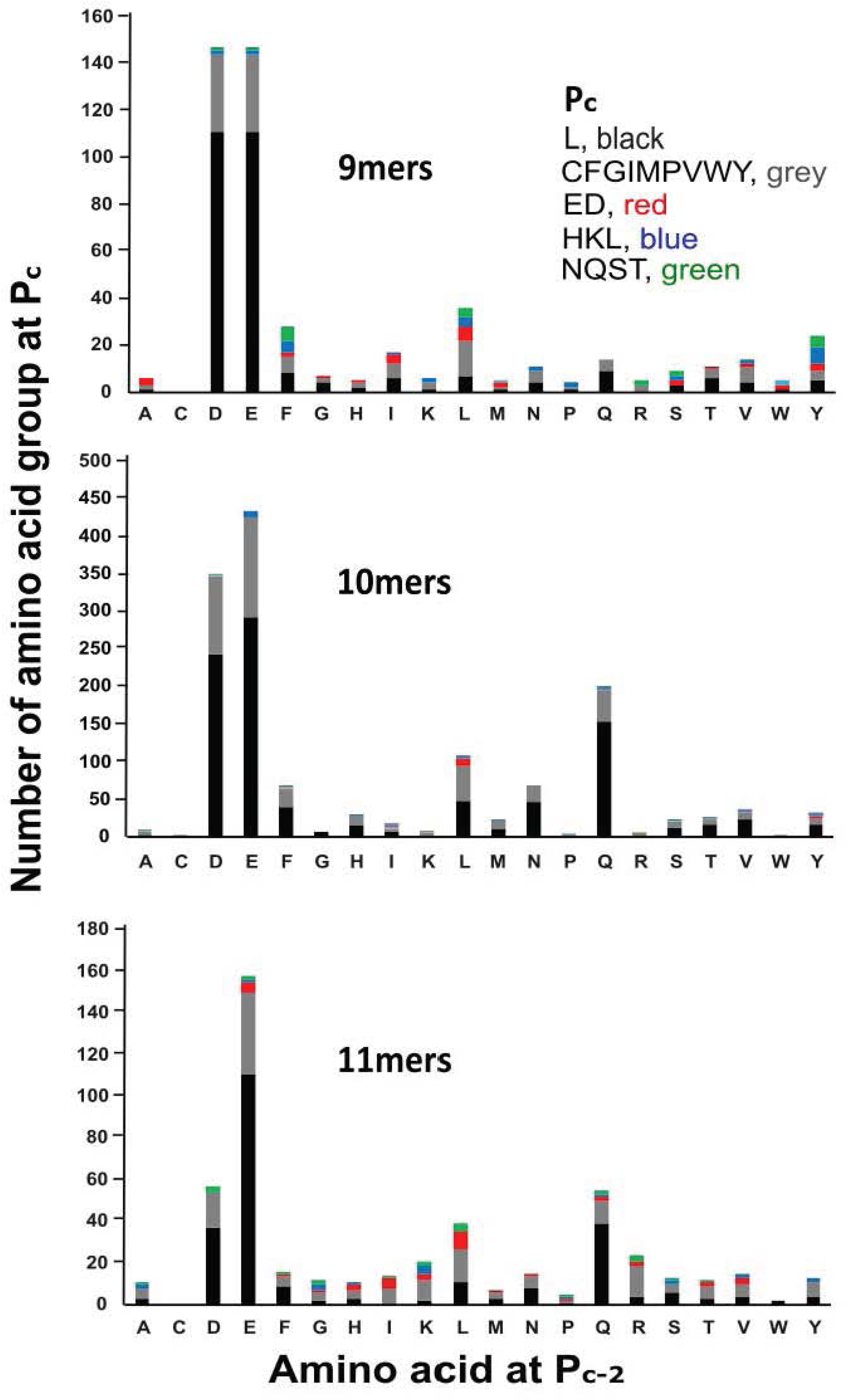
Superimposition of peptides from eight structures of BF2*21:01 show that the C-terminal portions all have similar conformations, while the N-terminal portions vary considerably. Bacterially-expressed heavy chain and β_2_-microglobulin were refolded in vitro with various synthetic peptides, purified by SEC, crystallized and the X-ray structures determined. Depicted are the main chains, with N-termini on the left, viewed from side of α2 helix of the class I molecule. Structures are 3BEV (GHAEEYGAETL, green) and 3BEV (REVDEQLLSV, light blue) (Koch et al., 2008), 2YEZ (TNPESKVFYL, pink), 4D0B (TAGQEDYDRL, yellow), 4D0C (TAGQSNYDRL, wheat) and 4CVZ (YELDEKFDRL, grey) (Chappell et al., 2015), and 5AD0 (GHAEEYGADTL, dark blue) and 5ACZ (GRAEEYGADTL, orange) (this publication).

One interpretation of this finding is that, during peptide binding, the C-terminus sits down first in the peptide-binding groove, with the N-terminus then accommodating as best it can. This idea fits with the importance of the C-terminus in peptide transport by the TAPs correlated with peptide binding to class I molecules (Deverson et al. 1998; Joly et al., 1998; Momberg et al., 1994) and with the action of ERAPs on the N-termini of bound but extended peptides (Li et al., 2019; Papakyriakou et al., 2018), but may not fit as well with other interpretations (Mavridis et al., 2021).

## Discussion

The dominantly-expressed class I molecule of the chicken MHC haplotype B21, BF2*21:01, remodels the binding site to bind very different peptides, with co-variation of the anchor residues at P_2_ and P_c-2_, so that no simple peptide-binding motif was obvious (Chappell et al., 2015; Koch et al., 2008). To better understand the rules of peptide binding, we used X-ray crystal structures, molecular modelling and thermostability assays, as well as assembly assays for single peptides, single-substitution libraries and double-substitution libraries assessed by SEC, by SEC followed by MALDI-TOF and by SEC followed by LC-MS/MS. In addition, we used immunopeptidomics (immunoaffinity chromatography followed by LC-MS/MS) of class I molecules from a B21 cell line. The overall conclusion of this study is that the rules of peptide binding to BF2*21:01 are complex, but a few simplifying principles contribute to the relatively simple overall outcome.

As previously inferred from structures of BF2*21:01 with six different peptides originally identified from a cell line, blood cells and spleen cells (Chappell et al., 2015; Koch et al., 2008), both the assembly assays and immunopeptidomics show co-variation of anchor residues at P_2_ and P_c-2_. The complexity of binding was enormous: out of 400 possible combinations at these two positions, 159 were found in double-substitution libraries of four peptide backbones assessed by SEC followed by LC-MS/MS and 260 were found by a single immunopeptidomics experiment with one cell line. Given that a single sampling will miss many rare sequences in immunopeptidomics (discussed below), it seems likely that most or all combinations of amino acids at P_2_ and P_c-2_ will turn out to be possible. This large number of combinations at P_2_ and P_c-2_ underscores the plasticity of binding allowed by BF2*21:01 remodelling the peptide-binding site.

However, it appears that the stability of binding is affected by length of peptide. Based on assembly assays, thermostability and number of peptides found by immunopeptidomics, 10mers appear to be the optimal length, with 9mers and 11mers tolerated if optimal amino acids at P_2_, P_c-2_ and P_c_ are present. There were some longer peptides found, but at much lower frequencies; these may bulge in the middle or hang out the C-terminal end (Burrows et al., 2006; Liu et al., 2012; Xiao et al., 2018).

The stability of binding is affected also by the particular residues present at P_2_ and P_c-2_, including their combinations. Not all amino acids are optimal, with His at P_2_ in the vast majority of 11mers, as assessed by immunopeptidomics. By assembly assay of peptide libraries, even the extremely stable 10mer REVDEQLLSV prefers His at P_2_. The widest range at P_2_ found for 10mers, as though they are able to tolerate less optimal amino acids. Even for 10mers, about a third of sequences each were found for Glu, for His and for the polar amino acids Asn, Gln and Ser, with the remainder mostly hydrophobic. Peptides with acidic residues at P_2_ vastly preferred Gln at P_c-2_, while basic and polar residues at P_2_ vastly preferred Asp and Glu at P_c-2_. These preferences were reflected in the optimal combinations for P_2_ and P_c-2_ by immunopeptidomics, with the most frequent combination for 11mers accounting for only 12% of total sequences, and with the most combinations and the most even distribution for 10mers. Many of these combinations were shared among 9mers, 10mers and 11mers, as though such anchor residues confer the greatest stability at all three lengths.

In addition, the amino acids present at P_3_ and P_c-3_ affect the anchor residues present at P_2_ and P_c-2_, further reducing the number of peptides that could be bound to BF2*21:01. The presence of particular residues at P_3_ affects the stability of peptide, as assessed by assembly and thermostability assays. Similarly, the combinations of P_2_ and P_3_ or of P_c-2_ and P_c-3_ are quite skewed as assessed by immunopeptidomics.

Finally, several amino acids are found at the anchor residue P_c_ by assembly assays with single peptides and with single-substitution libraries, but by immunopeptidomics far and away the most frequent was Leu. Assembly assays suggest that Ala and Trp are not favoured at P_c_ in the context of certain single peptides; one 11mer isolated from cells that had Trp at P_c_ had low thermostability, which was much improved by substitution with Leu. Assembly assays with peptide libraries based on the original 10mer and 11mer show that Leu/Ile, Phe, Met and Val are favoured. The polymorphic TAPs in chickens determine the peptides supplied to the class I molecules (Tregaskes et al., 2016; Walker et al., 2011), so it seems likely that the overwhelming Leu at P_c_ by immunopeptidomics is imposed by the translocation specificity in B21 cells, although translocation assays with single synthetic peptides show that other residues can be accommodated (Tregaskes et al., 2016). Whatever the reason, it is a further simplification reducing the number of peptides that can be bound to BF2*21:01 in vivo.

Confidence in the details of these interpretations must be tempered by the fact that both the assembly assays and immunopeptidomics have weaknesses, and that the two approaches do not give entirely the same results. Both MALDI-TOF and LC-MS/MS suffer from the weakness that peptides fly with different efficiencies, varying over orders of magnitude (Abdul-Khalek et al., 2025). In that sense, the peak height, ion current and number of reads are not quantitative representations of the amount of a given peptide present. For the assembly assays, it seems very likely the results for any particular position or combination of position will depend on the other amino acids present, the “backbone” of the particular peptide, so a wide range of peptide backbones must be assessed. Moreover, assembly assays assess the ability of a peptide to refold in vitro, which can differ significantly from the immunopeptidomics results from cells for which various quality control mechanisms are important. It is already known for another chicken class I allele, BF2*15:01, that refolding in vitro allows amino acids at anchor residues that are never found on the surface of cells by gas phase sequencing (Tregaskes et al., 2016; Wallny et al., 2006). Neither of these problems are so important for immunopeptidomics that can sample thousands of peptides simultaneously, but only a portion of very complex mixtures is sampled by the mass spectrometer, so both abundant and some of the rare peptides will be detected in each analysis. Therefore, the best way to determine whether a particular peptide is abundant is to detect it in many technical or biological replicates (for instance, Gastaldello et al., 2021; Halabi et al., 2021). In addition, the potential for contribution by the poorly-expressed class I molecule, BF1*21:01, to the immunopeptidomic analysis is a concern, although BF1 molecules are generally expressed at very low levels compared to BF2 molecules (Wallny et al., 2006). Also, only one biological replicate (with at least two technical replicates using different mass spec instruments) of a single cell line was assessed, and details may differ between cell types. The fact that results by the two approaches differed for P_c_ inspires further caution, particularly whether assembly assays with random libraries (for example, Jia et al., 2024; Li et al., 2024; Ma et al., 2020; Qu et al., 2019; Wei et al., 2021) accurately reflect biological situations (Tregaskes et al., 2016). Finally, chicken class I molecules allow peptides to extend out the C-terminal end of the peptide-binding groove (Xiao et al., 2018), so the assignment of P_c_ (and therefore P_c-2_ and P_c-3_) are not completely certain. The percentages of such C-terminal extensions are low for class I molecules with more easily understood peptide motifs (Han et al., 2023; Wallny et al., 2006), but only experimental approaches like crystal structures can resolve this issue for sure.

These concerns about the details of the interpretation aside, we have determined much about peptide-binding to the dominantly-expressed class I molecule of the B21 haplotype, an MHC haplotype associated with resistance to a variety of economically-important diseases. By in vitro assembly assays and from living cells, BF2*21:01 has the capacity to bind most combinations of the two anchor residues P_2_ and P_c-2_, making it by far the most promiscuous class I molecule so far described. However, the most frequent and stable peptides have strong preferences for length, and for identity of the amino acids found at P_2_ and P_c-2_, as well as backbone positions such as P_3_ and P_c-3_. In addition, although four amino acids at P_c_ allow assembly in vitro, only Leu at P_c_ is found with high frequency from the cell line. Some of these preferences can be understood by the structures and models determined in the course of this work. Taking these simplifying principles into account would allow prediction of peptides that will bind BF2*21:01 to better understand responses to infectious pathogens and to develop better poultry vaccines, but machine learning by artificial neural networks (Aranha et al., 2020; Jørgensen et al., 2014; Soam et al., 2011) may allow even better predictions. Having said that, at the moment predictions by machine learning can give answers, but without true knowledge, which is the understanding of why. The results and interpretations in this study are the first step towards that understanding.

## Materials and Methods

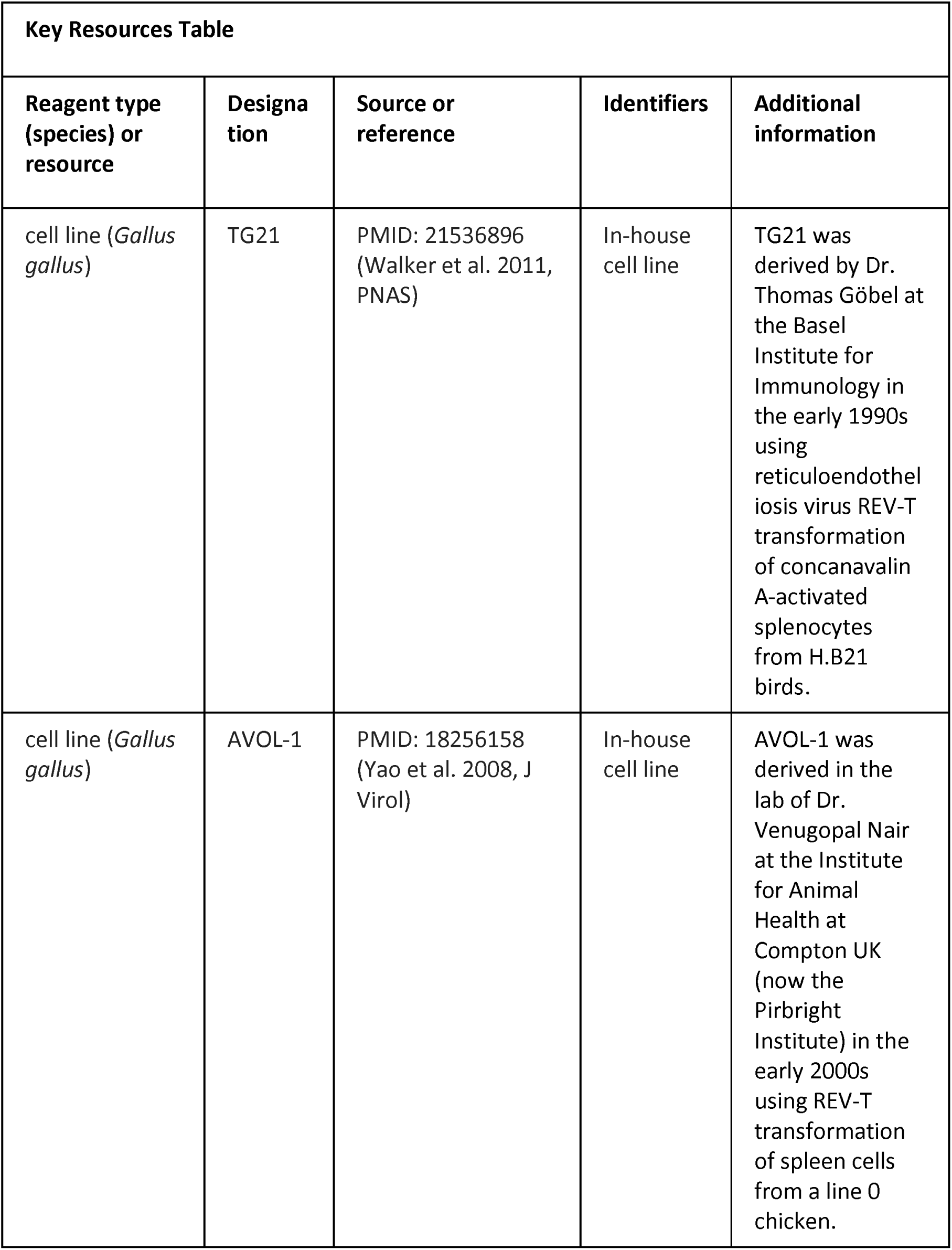
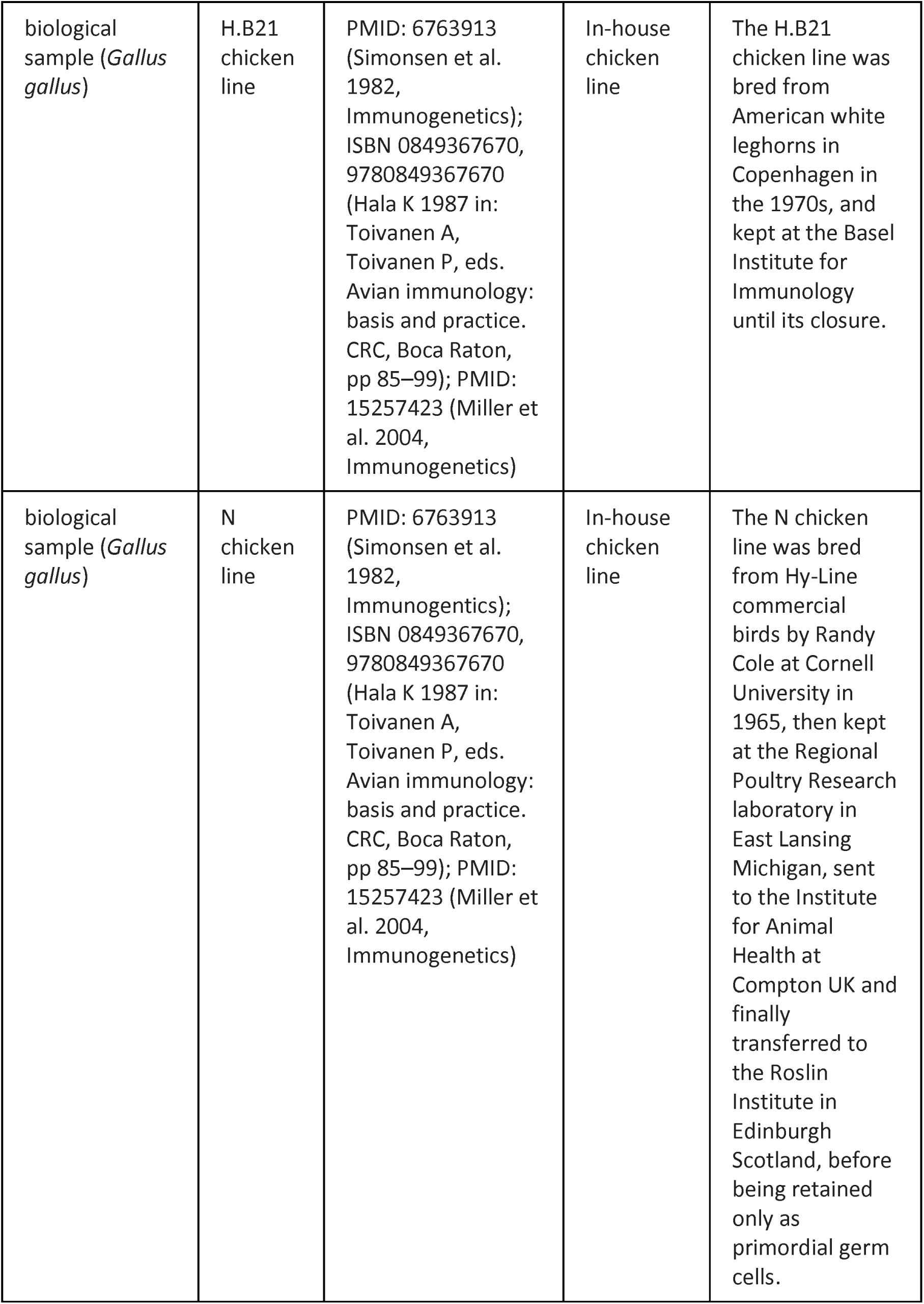
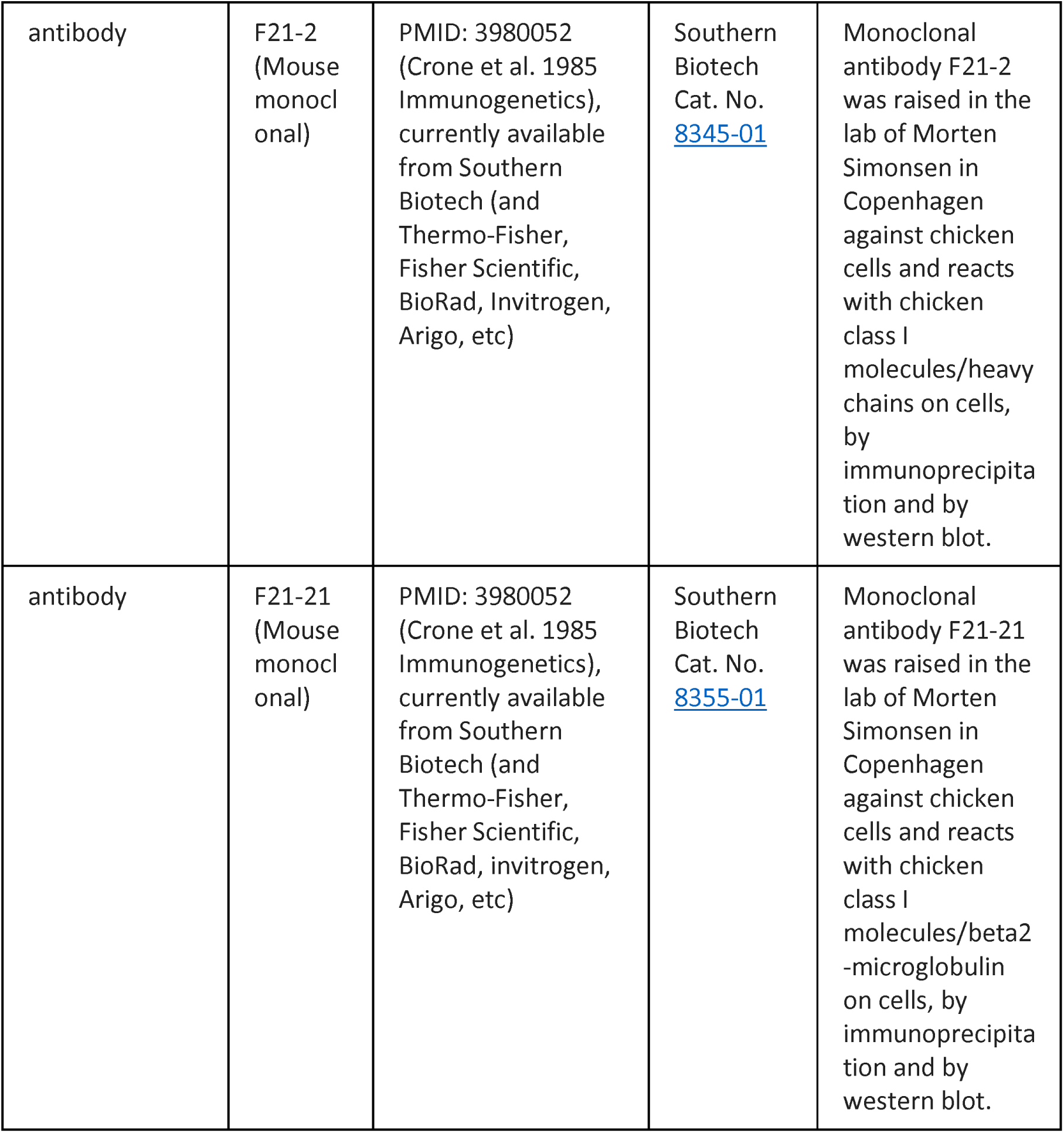

| Key Resources Table |  |  |  |  |
| --- | --- | --- | --- | --- |
| Reagent type (species) or resource | Designation | Source or reference | Identifiers | Additional information |
| cell line (*Gallus gallus*) | TG21 | PMID: 21536896 (Walker et al. 2011, PNAS) | In-house cell line | TG21 was derived by Dr. Thomas Göbel at the Basel Institute for Immunology in the early 1990s using reticuloendotheliosis virus REV-T transformation of concanavalin A-activated splenocytes from H.B21 birds. |
| cell line (*Gallus gallus*) | AVOL-1 | PMID: 18256158 (Yao et al. 2008, J Virol) | In-house cell line | AVOL-1 was derived in the lab of Dr. Venugopal Nair at the Institute for Animal Health at Compton UK (now the Pirbright Institute) in the early 2000s using REV-T transformation of spleen cells from a line 0 chicken. |
| biological sample (*Gallus gallus*) | H.B21 chicken line | PMID: 6763913 (Simonsen et al. 1982, Immunogenetics); ISBN 0849367670, 9780849367670 (Hala K 1987 in: Toivanen A, Toivanen P, eds. Avian immunology: basis and practice. CRC, Boca Raton, pp 85–99); PMID: 15257423 (Miller et al. 2004, Immunogenetics) | In-house chicken line | The H.B21 chicken line was bred from American white leghorns in Copenhagen in the 1970s, and kept at the Basel Institute for Immunology until its closure. |
| biological sample (*Gallus gallus*) | N chicken line | PMID: 6763913 (Simonsen et al. 1982, Immunogenetics); ISBN 0849367670, 9780849367670 (Hala K 1987 in: Toivanen A, Toivanen P, eds. Avian immunology: basis and practice. CRC, Boca Raton, pp 85–99); PMID: 15257423 (Miller et al. 2004, Immunogenetics) | In-house chicken line | The N chicken line was bred from Hy-Line commercial birds by Randy Cole at Cornell University in 1965, then kept at the Regional Poultry Research laboratory in East Lansing Michigan, sent to the Institute for Animal Health at Compton UK and finally transferred to the Roslin Institute in Edinburgh Scotland, before being retained only as primordial germ cells. |
| antibody | F21-2<br>(Mouse monoclonal) | PMID: 3980052<br>(Crone et al. 1985 Immunogenetics), currently available from Southern Biotech (and Thermo-Fisher, Fisher Scientific, BioRad, Invitrogen, Arigo, etc) | Southern Biotech Cat. No. 8345-01 | Monoclonal antibody F21-2 was raised in the lab of Morten Simonsen in Copenhagen against chicken cells and reacts with chicken class I molecules/heavy chains on cells, by immunoprecipitation and by western blot. |
| antibody | F21-21<br>(Mouse monoclonal) | PMID: 3980052<br>(Crone et al. 1985 Immunogenetics), currently available from Southern Biotech (and Thermo-Fisher, Fisher Scientific, BioRad, invitrogen, Arigo, etc) | Southern Biotech Cat. No. 8355-01 | Monoclonal antibody F21-21 was raised in the lab of Morten Simonsen in Copenhagen against chicken cells and reacts with chicken class I molecules/beta2-microglobulin on cells, by immunoprecipitation and by western blot. |

Cells and peptides. Blood was taken from H.B21 birds at the Gipf (Oberfrick) Farm of the Basel Institute for Immunology and processed to give peptides from class I molecules as described (Wallny et al., 2006). Spleens were taken from N line birds at the Institute for Animal Health (Compton, UK) after retirement from breeding flock and schedule 1 procedures, and processed to give peptides from class I molecules as described (Chappell et al., 2015). The B21 cell line TG21 was derived by reticuloendotheliosis virus REV-T transformation of concanavalin A-activated splenocytes from H.B21 birds at the Basel Institute for Immunology and maintained as described (Walker et al., 2011), with peptides isolated from class I molecules as described (Chappell et al., 2015). The peptides from blood, spleen and TG12 cells were separated by HPLC and single peaks subjected to gas phase sequencing as described (Chappell et al., 2015; Wallny et al., 2006). The B21 cell line AVOL-1 was derived by REV-T transformation of spleen cells from a line 0 chicken at the Pirbright Institute, with peptides isolated from class I molecules and prepared for immunopeptidomics, all as described (Chappell et al., 2015).

Expression and isolation of protein chains. The cloning of BF2*21:01 and β_2_-microglobulin sequences, recloning into pET22b(+) expression plasmids, expression in BL21 (λDE3) pLysS Rosetta cells, isolation, washing and solubilisation of inclusion bodies using urea were broadly as previously described (Chappell et al., 2015; Koch et al., 2007), but occasionally used TopTen medium, which led to much higher yields.

Refolding with peptides. Renaturation of BF2*21:01 heavy chain with β_2_-microglobulin and synthetic peptides, separation by size exclusion chromatography and concentration of monomer fractions were broadly as previously described (Chappell et al., 2015; Koch et al., 2007). Single peptides and peptide libraries were synthesised by fmoc chemistry and appropriately purified (Pepceuticals), with lyophilised material dissolved in 0.1 % trifluoroacetic acid. The peptide libraries had one or two positions at which all natural amino acids except cysteine were coupled, using ratios of reactants previously determined to give more-or-less equivalent incorporation. The renaturation buffer used was 400 mM arginine, 2 mM ethylene diamine tetraacetic acid (EDTA), 100 mM TrisCl pH 8.2, with oxidized glutathione added to 0.5 mM, reduced glutathione added to 5 mM and 4-(2-aminoethyl)benzenesulfonyl fluoride hydrochloride (AEBSF) added to 0.5 mM just before use. The refolding involved adding denatured heavy chain with rapid stirring to refolded β_2_-microglobulin and synthetic peptide(s) in renaturation buffer, and incubating with continued stirring 18-40 hours at 4°C. The samples were centrifuged at high speed to precipitate flocculate material, and the supernatant was passed through 0.2 or 0.45 micrometer sterile filters before separation by size exclusion chromatography in TBS (100 mM NaCl, 25 mM TrisCl pH 8.0) using an AKTA 920 running at 1 ml/min at room temperature, followed by concentration of fractions corresponding to refolded monomers.

The total volume of refold mixture was typically 100 ml for single peptide refolds intended for crystallization and for thermostability assays, and 1 ml or 1.8 ml for all other refolds involving peptide libraries. Most refolding was at molar concentrations of roughly 1 mole heavy chain: 2 mole β_2_-microglobulin: 10 mole peptide (1:2:10), except for some analytical runs with peptide libraries in which 1:2:10 was compared with 1:2:2 (the latter to control for one peptide outcompeting other peptides with slightly lower affinity). Heavy chain was added slowly dropwise to rapidly stirred refold mix containing peptide and previously refolded β_2_-microglobulin. After refolding at 4°C, some samples were analysed immediately, while others were incubated at 4°C or at 42°C (the latter to assess whether the complexes stable at 4°C might unfold at the body temperature typical of a chicken) for 1, 6 or 18 hours (results shown for 18 hours in Fig. 8).

The larger scale refolds were chromatographed on a HiLoad 26/60 Superdex 200 column (GE Healthcare, UK) column at room temperature. Fractions corresponding to monomer were pooled and concentrated with a Vivaspin 50 (10 kDa cut-off, Sartorius) at 4°C until the volume was a few ml. Then, a Vivaspin 2 (10 kDa cut-off, Sartorius) at 4°C was used to replace the buffer with 50 mM NaCl, 10 mM TrisCl pH 8 and to reduce the volume to 100-200 microliters, which were snap frozen on dry ice before storage at −80°C until use.

The smaller scale refolds were chromatographed on a Superdex 200 10/300 GL column (GE Healthcare, UK) column, with fractions corresponding to monomer pooled and concentrated first using a pre-rinsed Amicon Ultra-4 10 kDa column and then a Vivaspin 500 10 kDa column to 100-200 microliters, and the buffer exchanged to 10 mM TrisCl pH 8. The sample was further concentrated using a SpeedVac at 40°C to 5-10 microliters, glacial acetic acid was added to 5% v/v followed by further concentration by SpeedVac at 40°C to 2-4 microliters, before freezing at −80°C. Analysis was typically by matrix-assisted laser-desorption ionisation-time of flight mass spectroscopy (MALDI-TOF, Protein and Nucleic Acid Chemistry services, Department of Biochemistry, University of Cambridge), but later by liquid chromatography-mass spectroscopy/mass spectroscopy (LC-MS/MS, Cambridge Centre for Proteomics, Department of Biochemistry, University of Cambridge). For MALDI-TOF, spectra were analysed by comparison to mass-to-charge (m/z) ratio calculated for each peptide in the library, with the amount of each peptide detected in the input mixture verified by MALDI-TOF. For LC-MS/MS, the peptides were identified using Mascot. Calculations, comparisons and graph-making were performed using Excel, and figures made using Inkscape.

Thermostability assays. Refolds of 6 ml were performed and split into 1 ml aliquots at 1 mg/ml. The thermal stabilities of BF2*21:01 with different peptides were assessed using the ThermoFluor assay (Matulis et al., 2005). In detail, 20 μL of each protein solution at 1 mg/mL was added to 5 μL of SPYRO orange dye (Life Technologies) (1000-fold dilution) in a 96-well plate. The temperature was increased at a rate of 1.5°C per minute and fluorescence monitored using a Stratagene MX3005P real time PCR machine. Fluorescence measurements were normalised and then fitted to a Boltzmann sigmoidal curve using GraphPad Prism to calculate melting temperatures.

Crystallography. Protein crystals were obtained and structures were determined overall as described (Chappell et al., 2015). In detail, recombinant MHC class I complexes were crystallized using the sitting-drop method [0.2 M MgCl_2_.(H_2_O)_6_, 0.1 M Tris, pH 8.0, 20% PEG6000 for 5AD0 with peptide GHAEEYGADTL; Molecular Dimensions E9, pH 8.5 for 5ACZ with peptide GRAEEYGADTL]. Crystals were flash frozen in liquid nitrogen and native data sets for each crystal were collected at 100K [ESRF, Grenoble (beamline ID23-2) for 5AD0; Diamond Light Source, Harwell (beamline I02) for 5ACZ]. Data were processed using either AUTOPROC (Vonrhein et al., 2011) or XIA2 (Winter, 2010) with XDS (Kabsch, 2010) for integration and AIMLESS (Evans, 2006) or SCALA (Evans and McCoy, 2008) for scaling. All structures were solved by molecular replacement, as implemented in Phaser (McCoy et al., 2007), part of the CCP4 software package (Winn et al., 2011). Starting molecular replacement models were generated using CHAINSAW (Stein, 2008) and the atomic co-ordinates of the chicken B21 MHC class I molecule (PDBID: 3BEV) with the peptide removed. Model building and refinement were carried out using COOT (Emsley et al., 2010) and AUTOBUSTER (Bricogne et al., 2011) or REFMAC (Murshudov et al., 1997), with the heavy and light chains of the MHC molecule rebuilt first before the peptide was modelled into residual electron density. Structures were depicted using PyMol (The PyMOL Molecular Graphics System, Version 2.0 Schrl1dinger, LLC).

Modelling. Beginning with known crystal structures, modelling was carried out using PyMol with the protein mutagenesis Wizard, the rotamer toggle, show bumps and show surfaces. Immunopeptidomics. Peptide sequences from AVOL-1 cells were determined just as described (Chappell et al., 2015). In detail, immunoaffinity beads were produced, with all steps at room temperature. Protein G-Sepharose beads (Expedeon) were washed with 50 mM borate, 50 mM KCl, pH 8.0, the equivalent of 1-ml packed beads was incubated with 3 mg F21-2 (produced by the Microbiological Media Services of the Pirbright Institute) for 1 hr, treated with 40 mM dimethyl pimelimidate dihydrochloride (Sigma) in 0.1 M triethanolamine, pH 8.3 for 1 hr to cross-link the antibody to the protein G, washed with 100 mM citric acid pH 3.0, and equilibrated in 50 mM TrisHCl, pH 8.0. The cell line AVOL-1 was washed with PBS. A pellet of 10^9^ cells was incubated with 10 ml 1% Igepal 630, 300 mM NaCl, 100 mM Tris pH 8.0 for 30 min at 4°C, subcellular debris was pelleted by centrifugation at 300×g for 10 min and 15,000×g for 30 min at 4°C, and the cleared lysates were incubated with 1 ml immunoaffinity beads for 1 hr at 4°C. The beads were washed with 50 mM TrisCl, pH 8.0 buffer, first with 150 mM NaCl, then with 400 mM NaCl and finally with no salt.

Bound material was eluted with 10% acetic acid. The eluted material was dried, resuspended in 3% acetonitrile, 0.1% formic acid in water, loaded directly onto on a 4.6 × 50 mm ProSwiftTM RP-1S column (ThermoFisher) and eluted at 500 μl/min flow rate for 10 min with a linear gradient from 2 to 35% buffer B (0.1% formic acid in acetonitrile) in buffer A (0.1% formic acid in water) using an Ultimate 3000 HPLC system (ThermoFisher), with fractions collected from 2 to 15 min. Protein detection was performed at 280 nm absorbance, with fractions eluting before β_2_-microglobulin pooled and dried.

For liquid chromatography tandem mass spectrometry (LC-MS/MS), peptides were analysed using either a Q-Exactive (Thermo Scientific) or a TripleTOF 5600 (AB Sciex) system, as identified (Dataset 1). For the QExactive system, peptides were separated on a Ultimate 3000 RSLCnano System utilizing a PepMap C18 column, 2 μm particle size, 75 μm × 50 cm (Thermo Scientific) with a linear gradient from 3% to 35% buffer B in buffer A (as above) at a flow rate of 250 nl/min (∼ 65 MPa) for 60 min, and the 15 most intense precursors per full MS scan were selected for MS/MS analysis using HCD fragmentation. For the TripleTOF system, peptides were separated with a 15 cm × 75 μm ChromXP C18-CL (3 μm particle size) using an ekspert nanoLC 400 cHiPLC system (Eksigent) with a linear gradient from 8% to 35% buffer B in buffer A (as above) at a flow rate of 300 nl/min (∼1600 psi) for 60 min, and CID fragmentation was induced on the 30 most abundant ions per full MS scan. All fragmented precursor ions were actively excluded from repeated selection for 15 s. Data were analysed using Peaks 7 (Bioinformatics Solutions) using a database containing all 24,092 Uniprot entries for the organism Gallus gallus combined with protein translations (>8 amino acids) of either all six reading frames of gallid herpesvirus 2 (NCBI entry NC_002229.3; 10,026 entries) or reticuloendotheliosis virus (NC_006934.1; 412 entries). Results were filtered using a false discovery rate of 1% that was determined by parallel searching of a randomized decoy database. Peptide motifs were determined by Gibbs clustering (Andreatta et al., 2013) through a website (https://services.healthtech.dtu.dk/services/GibbsCluster-2.0/).

## Acknowledgements

many thanks to researchers for help to identify the first peptides (Fig. 1) at the Basel Institute for Immunology including Jan Salomonsen, and at the IAH Compton including Lawrence Hunt, Fiona Johnston, Denise A. Marston, Andy van Hateren, Simon Camp, Jansen P. Jacob, Iain Shaw, Timothy J. Powell, Sally Rogers and Brian Walker. Also thanks to Pietro Roversi at Oxford for help with the structures, and Laura Mears, Xaquin Castro-Dopico and Len Packman for help with the peptide libraries. Funding at the time the research was being done included support from Hofmann La Roche at the Basel Institute for Immunology to JS, DWA, HJW and JK, and from the Biotechnology and Biological Sciences Research Council (BBSRC) at the Institute for Animal Health Compton (later renamed the Pirbright Institute) to VN, WM and JK. In addition, grants included Welcome Trust Studentships to LM and CH in Cambridge and a BBSRC studentship to PC at Oxford, Wellcome Trust Senior Investigator Award to SML at Oxford, grants to NT at Oxford, a program grant (089305/Z/09/Z) and an Investigator Award from the Wellcome Trust (110106/Z/15/Z) to JK in Cambridge and later in Edinburgh, and a BBSRC project grant (BB/V000756/1) and a German Science Foundation (DFG) consortium grant (FOR5130 - ImmunoChick, KA 5564/1) to JK in Edinburgh.

## Author contributions

Conception and design: VN, SML, NT, JK

Acquisition of data: MH, PEC, MJD, ŁM, CH, MJD, HJW, DWA, WM

Analysis and interpretation of data: MH, PEC, SH, MJD, KSL, HJW, DWA, WM, NT, SML, JK

Drafting or revising the article: EMM, SH, NT, SML, JK (all authors had the opportunity to read and comment on the article)

Ethics involving chickens were carried out at the Basel Institute under the provisions in place at the time, and at the Institute for Animal Health under Home Office project license PPL 80/2420 and with ethical approval of the Local Ethical Review Committee.

Fig. 11 – Source Data 1. Raw data for ion current (counts) from MALDI-TOF and LC-MS/MS analyses of the peptides that assembled with BF2*21:01 from six double-substitution libraries (source of data for bar graphs in Figs. 11-13).

Fig. 17 - Source Data 1. Raw data from immunopeptidomics of class I molecules from AVOL-1 cells (source of data for Figs. 17-27).

---

**Table 1.**
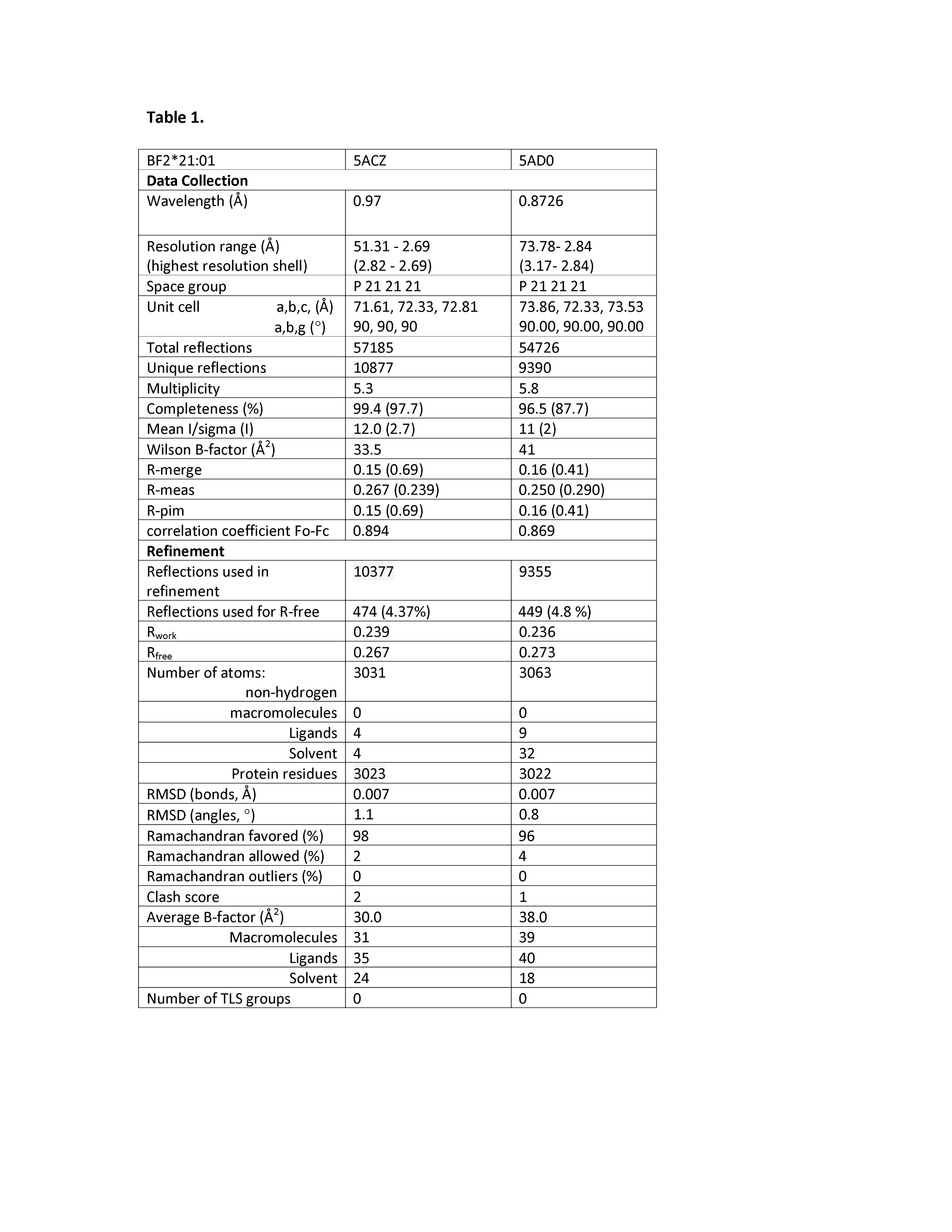

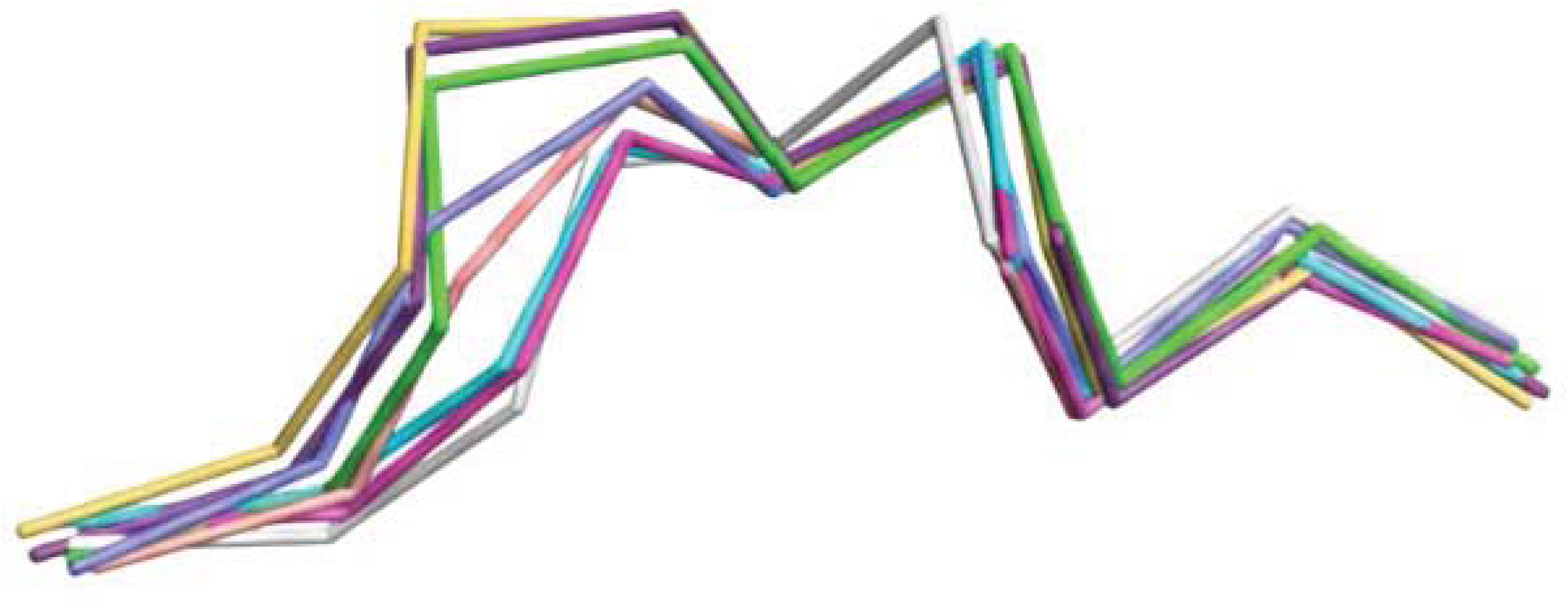

